# Nucleoporin Elys attaches peripheral chromatin to the nuclear pores in interphase nuclei

**DOI:** 10.1101/2023.08.16.553518

**Authors:** Semen A. Doronin, Artem A. Ilyin, Anna D. Kononkova, Mikhail A. Solovyev, Oxana M. Olenkina, Valentina V. Nenasheva, Elena A. Mikhaleva, Sergey A. Lavrov, Anna Y. Ivannikova, Anna A. Fedotova, Ekaterina E. Khrameeva, Sergey V. Ulianov, Sergey V. Razin, Yuri Y. Shevelyov

## Abstract

Transport of macromolecules through the nuclear envelope (NE) is mediated by nuclear pore complexes (NPCs) consisting of nucleoporins (Nups). Elys/Mel-28 is the Nup that binds and connects the decondensing chromatin with the reassembled NPCs at the end of mitosis. Whether Elys links chromatin with the NE during interphase is unknown. Using DamID-seq, we identified Elys binding sites in *Drosophila* late embryos and divided them into those associated with nucleoplasmic or with NPC-linked Elys. These Elys binding sites are located within active or inactive chromatin, respectively. Strikingly, *Elys* knockdown in S2 cells results in peripheral chromatin displacement from the NE, in decondensation of NE-attached chromatin, and in derepression of genes within. It also leads to slightly more compact active chromatin regions. Our findings indicate that NPC-linked Elys, together with the nuclear lamina, anchors peripheral chromatin to the NE, whereas nucleoplasmic Elys decompacts active chromatin.

**Author summary:** Heterochromatin in interphase nucleus is localized mostly at the nuclear periphery. However, the forces maintaining its peripheral localization are not well understood. Nuclear envelope consists of two lipid bilayer membranes separated by perinuclear space. The inner nuclear membrane is lined by the nuclear lamina, and both membranes are pierced by nuclear pore complexes composed of nucleoporins. Nuclear envelope can serve as a scaffold to which heterochromatin is attached. In the present study, we identified nucleoporin Elys as one of the key players maintaining peripheral localization of heterochromatin during interphase. Elys binds to multiple genomic sites located within heterochromatin and thus links it to nuclear pore complexes. However, the nucleoplasmic fraction of Elys binds to active genes and enhancers, resulting in decompactization of their chromatin.

## Introduction

Although the idea that chromosomes in interphase nucleus are attached to the nuclear envelope (NE) was proposed a long ago [1], mechanisms mediating this attachment are still poorly understood. Accumulating evidence suggests that peripheral chromatin is in intimate association with both the nuclear lamina (NL) and the nuclear pore complexes (NPCs). NL consists of Lamins and Lamin-associated proteins [2]. Genomic regions interacting with the NL (lamina-associated domains, LADs) were successfully identified in different organisms, including *Drosophila*, mammals, and nematode [3–8]. Depletion of specific components of the NL resulted in the displacement of chromatin from the NE to nuclear interior [9–19], thus indicating that chromatin is attached to the NL and that interphase chromosomes are slightly stretched by this attachment [19,20]. In addition, chromosomes from yeast, *Drosophila* and mammals interact with various nucleoporins (Nups) (i.e., proteins composing NPCs) [21–37]. However, contrarily to yeast, Nups-chromatin interactions in metazoans may take place not only at the NE, but also in the nuclear interior [27–29,31]. For example, in *Drosophila*, nucleoplasmic fraction of Nup98 interacts with active chromatin, whereas Nup98, as a component of NPCs, mostly associates with inactive chromatin [28].

Several findings point to the involvement of NPCs in chromatin tethering to the NE in metazoans. The dosage-compensated single X chromosome in *Drosophila* SL-2 cells is stronger bound with Nup153 and is localized closer to the NE than autosomes [29], whereas Nup153 depletion leads to repositioning from the NE of several Nup153-target loci in both *Drosophila* SL-2 and mouse embryonic stem cells [29,33]. Similarly, Nup Elys/Mel-28 (further Elys) interacts stronger with *C*. *elegans* male single X chromosome than with two X chromosomes of hermaphrodites which correlates with less distant position of the male X chromosome from the NE [38]. In addition, Nup155 is involved in chromatin tethering to the NE in *Drosophila* S2 and ovarian germline cells [39]. However, despite these indications, the contribution of NPC-chromatin interactions to the maintenance of peripheral chromatin positioning as well as an impact of these interactions on the overall genome architecture and gene expression remain largely unexplored.

Elys may be the unique Nup that directly tethers chromatin to the NE [40]. Initially, it was identified as the AT-hook-containing transcription factor expressed in embryonic haematopoietic tissues of mouse [41]. More recent experiments have shown that, during mitosis, Elys, together with the Nup107-160 complex, are concentrated on kinetochores in mammals, *Xenopus* and the nematode [42–45]. Importantly, upon mitotic exit, Elys participates in the NPC reassembly by connecting the Nup170-160 complex to decondensing chromatin [42,45–47]. Accordingly, in the nematode, *Xenopus* and mammalian cells, Elys depletion results in the partial loss of nuclear pores from the NE [42,44,45,48]. Partial loss is explained by an existence of Elys-independent pathway of NPC incorporation into the NE during interphase [48]. Knockdown of *Elys* in *Drosophila* salivary glands caused the enhanced apoptosis and the disappearance of Lamins and Nups from the NE [49].

Eight Elys molecules, as the components of NPC nuclear basket, face peripheral chromatin [50]. Elys contains β-propeller and α-helical domains at the N-terminal part, which are responsible for its association with the Nup107-160 complex [51,52]. The C-terminal part of Elys bears AT-hook DNA-binding motif recognizing A/T-rich DNA sequences [47] and an additional domain, able to bind with the acidic patch of a nucleosome [52–56]. *Drosophila* Elys also possesses β-propeller and α-helical domains, but contains three non-canonical AT-hook-like motifs which were shown to be required for Elys binding to A/T-rich sequences *in vitro* [49].

Elys interacts with SWI/SNF chromatin remodeling complex PBAP [36] known to remove nucleosomes from the regulatory regions of genome and to switch chromatin to a more open state [57–59]. Consistent with this ability, artificial tethering of Sec13, the partner of Elys in the Nup107-160 complex, to several sites on *Drosophila* polytene chromosomes resulted in the recruitment of Elys to these sites followed by local chromatin decondensation [36]. Yet, ATAC-seq assay did not reveal notable changes in chromatin accessibility upon Elys depletion in S2 cells [60]. Elys binding sites across the entire *Drosophila* genome were recently identified by ChIP-seq in brains from 3^rd^ instar larvae and in embryonic S2 cell line [35,37]. Elys ChIP-seq profile appeared to be highly correlated with the profile of H3K27 acetylation [35]. At the same time, a fraction of ChIP-seq sites common between Elys and Nup93 or between Elys and Nup107 was found to reside within LADs, thus indicating peripheral localization of the chromatin carrying these sites [37]. However, Elys ChIP-seq sites were not explicitly classified as bound by nucleoplasmic or by NPC-linked Elys. Therefore, the influence of Elys on chromatin state and on gene expression specifically in these two different locations was unclear. Moreover, it was not previously explored whether Elys tethers chromatin to the NE and maintains its peripheral positioning during interphase.

Here, we report that knockdown of *Elys* (Elys-KD) in *Drosophila* S2 cells does not notably affect the abundance of NPCs at the NE and does not result in the increased apoptosis, thereby keeping cells alive. These findings allow to consider S2 cells as an appropriate model to analyze changes in genome architecture upon the lack of Elys. Using S2 cells and late embryos, we found that Elys, as a component of NPCs, binds with numerous genomic sites within LADs and this binding is required for the retention of peripheral chromatin at the NE during interphase. Consistent with these results, we found that, upon Elys-KD in S2 cells, the topologically associating domains (TADs [61–64]) that are attached to the NL become less condensed and genes located within them are slightly derepressed. These effects are similar to what was observed upon disruption of the NL in S2 cells [19]. Taken together, these findings support the model according to which the components of the NL, in cooperation with the NPC-linked Elys, bind peripheral chromatin and maintain its proper localization and functions in interphase nuclei.

## Results

### Elys-KD in *Drosophila* S2 cells does not lead to considerable loss of NPCs at the NE

Knockdown of *Elys* in salivary glands from *Drosophila* third instar larvae impairs localization of major NE components, such as Nups, Lamins, and Lamin-B-receptor (LBR) [49]. Since salivary glands contain non-dividing cells undergoing endoreplication, we examined whether Elys is similarly required for correct localization of NE components in the mitotically dividing embryonic S2 cells. We stained S2 cells with anti-Elys antibodies generated earlier [65] and revealed Elys localization mostly at the nuclear rim (Fig 1A, upper left panel). Elys colocalizes with the NPCs (stained with Mab414 antibodies which detect the phenylalanine-glycine-rich Nups) at the NE giving a characteristic punctate pattern (Fig 1A, upper right panel). RNAi knockdown of Lamin *Dm0* (Lam-KD; S1A Fig) in S2 cells results in NPC clustering [66] manifested in a more discrete staining of NPCs and their constituent Elys at the NE (Fig 1A, middle panel). However, contrary to the observations in salivary glands, Elys-KD in S2 cells resulting in 6-13 fold decrease of Elys protein level (S1 Fig) does not impair Lamin localization (Fig 1A, low left panel). Strikingly, unlike in nematode, *Xenopus*, and mammals [42–46], and unlike in *Drosophila* salivary glands [49], Elys-KD in S2 cells does not lead to a notable loss of NPCs at the NE and to the emergence of cells lacking nuclear pores (Fig 1A, low right panel, and Fig 1B and 1C). Only knockdown of both Lamin *Dm0* and *Elys* (Fig 1D, upper panel) results in the pronounced reduction of NPC staining at the nuclear rim (Fig 1D, lower panel).

**Fig 1.**
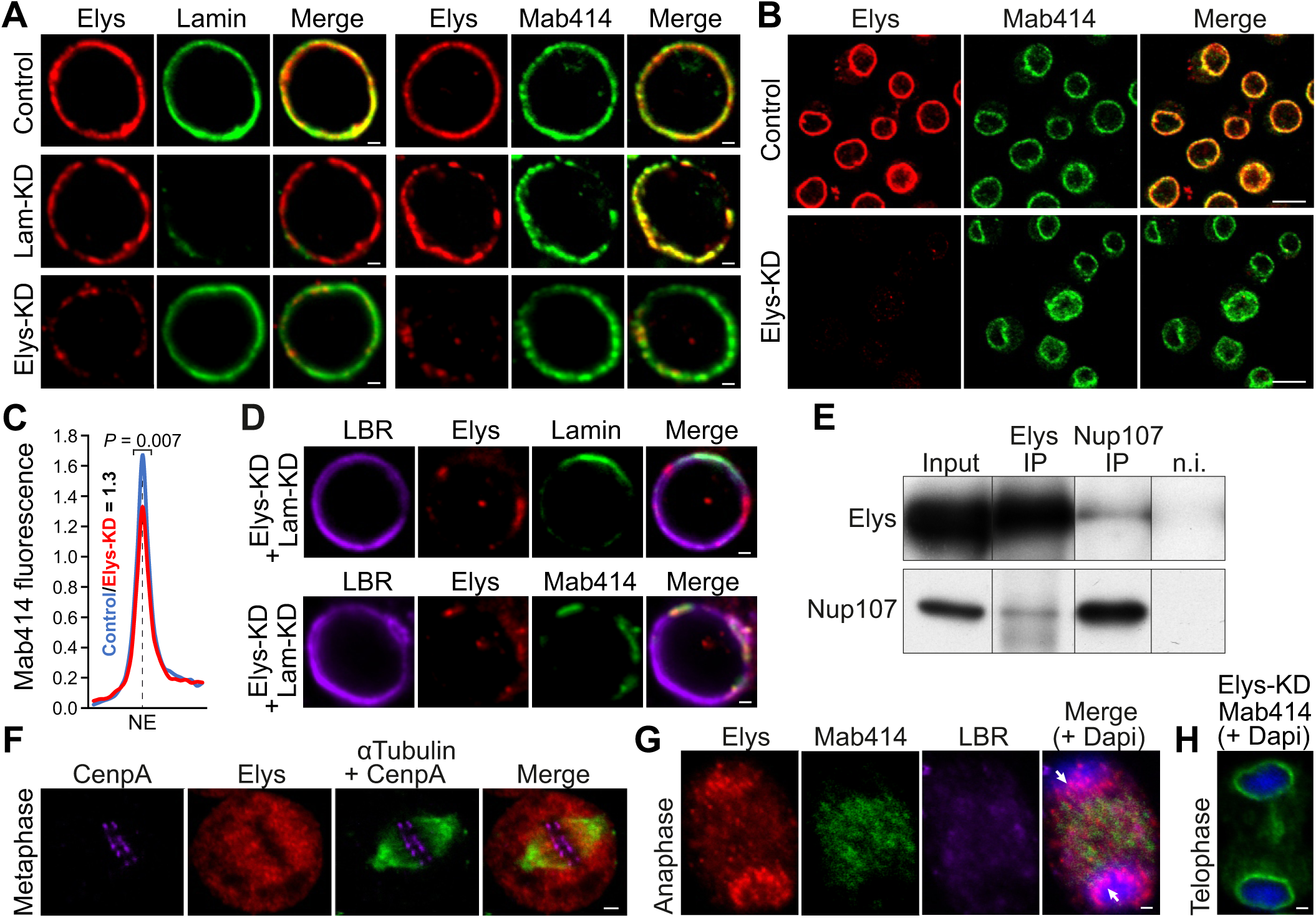
Elys-KD in S2 cells does not lead to a notable loss of NPCs at the NE. (**A**,**B**,**D**) Immunostaining of control, Lam-KD, Elys-KD, or (Lam-KD plus Elys-KD) S2 cells with anti-Elys, anti-Lam and Mab414 antibodies (**A**), with anti-Elys and Mab414 antibodies (**B**), or with anti-LBR, anti-Elys and anti-Lam antibodies (**D**). Scale bars 1 µm (**A**,**D**), 10 µm (**B**). (**C**) *ImageJ* quantification of Mab414 average fluorescence intensity (normalized on average Dapi fluorescence) across the NE in Elys-KD (two replicates, n = 75) and control (two replicates, n = 75) S2 cells. *P* value was estimated in a M-W *U*-test. (**E**) Western-blot analysis of proteins, co-immunoprecipitated with anti-Elys or anti-Nup107 antibodies from S2 extracts, probed by anti-Elys, or anti-Nup107 antibodies (n.i. – non-immune serum, IP/input ratio 1:4.5). (**F**,**G**) Immunostaining of S2 cells with anti-CenpA (kinetochores, violet), anti-Elys (red), anti-α-Tubulin (green) in metaphase (**F**), with anti-Elys (red), anti-Mab414 (green), anti-LBR (violet), Dapi (blue) in anaphase (**G**). Scale bars 1 µm (**F**,**G**). Arrows point to the Elys concentrated around decondensing chromatin (**G**). (**H**) Immunostaining of Elys-KD S2 cells with Mab414 antibodies (green) counterstained with Dapi (blue) in telophase. Scale bar 1 µm.

These surprising results prompted us to examine Elys behavior during mitosis in S2 cells. Post-mitotic NPC reassembly in other organisms is initiated in anaphase by Elys binding to the decondensing chromatin, followed by Elys-mediated recruitment of the Nup107–160 subcomplex of NPC [42,45–48,67,68]. We found that Elys and Nup107 reciprocally co-immunoprecipitate each other from protein extracts of S2 cells (Fig 1E), therefore indicating a physical association between these Nups, at least during the interphase. In mammals, *Xenopus* and nematode, a fraction of Elys, being in the complex with Nup107-160, is localized on kinetochores during mitosis [42–45,52]. However, Mehta *et al*. [49] revealed Elys not staying on kinetochores in *Drosophila*. We confirmed their findings by staining of S2 cells with anti-α-tubulin and anti-Elys antibodies, which demonstrates diffuse Elys distribution around condensed chromosomes, but not at the kinetochores during metaphase (Fig 1F). In anaphase, Elys appears to concentrate around decondensing chromatin earlier than other Nups or LBR (Fig 1G), thus pointing to the Elys leading role in determining seeding sites on chromatin upon the post-mitotic NPCs reassembly in *Drosophila* S2 cells, as shown in other organisms [42,45–48,67,68]. Nevertheless, the lack of Elys in S2 cells does not prevent the appearance of NPCs at the NE at the end of mitosis (Fig 1B). Moreover, upon Elys-KD, the incorporation of NPCs in the NE occurs when cytokinesis is not yet complete (Fig 1H). Consistent with these results, we did not reveal any disturbance of mRNA export from the nucleus upon Elys-KD (S2 Fig), thus indicating that nuclear-cytoplasmic traffic of RNA via nuclear pores functions normally. We conclude that Elys-dependent incorporation of NPCs in the NE at the end of mitosis may be bypassed in S2 cells by rapidly acting Elys-independent mechanism.

RNAi knockdown of *Elys* using ubiquitous *act5C*-GAL4 driver was shown to induce apoptosis in *Drosophila* salivary glands [49]. We explored whether Elys-KD in S2 cells also results in the enhanced cell death. Unexpectedly, TUNEL assay did not detect any increase in the proportion of apoptotic cells upon Elys-KD in comparison with the control cells (3.6% versus 3.3%, respectively, *P* = 0.99, M-W U-test, S3 Fig, S1 Table).

In summary, the absence of notable NPC and NL defects, as well as normal cell viability upon Elys-KD makes S2 cells an appropriate model to explore the influence of Elys depletion on the genome architecture.

### Elys binds to numerous sites on polytene chromosomes

Several Nups, including Sec13, occupy dozens of sites on polytene chromosomes mostly located away from the NE [27]. Artificial tethering of Sec13 to several ectopic sites on polytene chromosomes results in recruiting of Elys to these sites, correlated with local chromatin decondensation [36]. Using immunostaining of squashed polytene chromosomes with anti-Elys antibodies, we revealed that Elys by itself binds to the plethora of sites, some of them are colocalized with polytene chromosome bands (pink strips on Fig 2A), while others are colocalized with inter-bands (red strips on Fig 2A). Polytene chromosome bands and inter-bands are known to contain silent and expressed genes, respectively [69]. Since numerous fluorescence *in situ* hybridization (FISH) experiments indicate that active chromatin is located far from the NE [70], we propose that similarly to some other Nups, Elys interacts with chromatin positioned both at the NE and in the nuclear interior.

**Fig 2.**
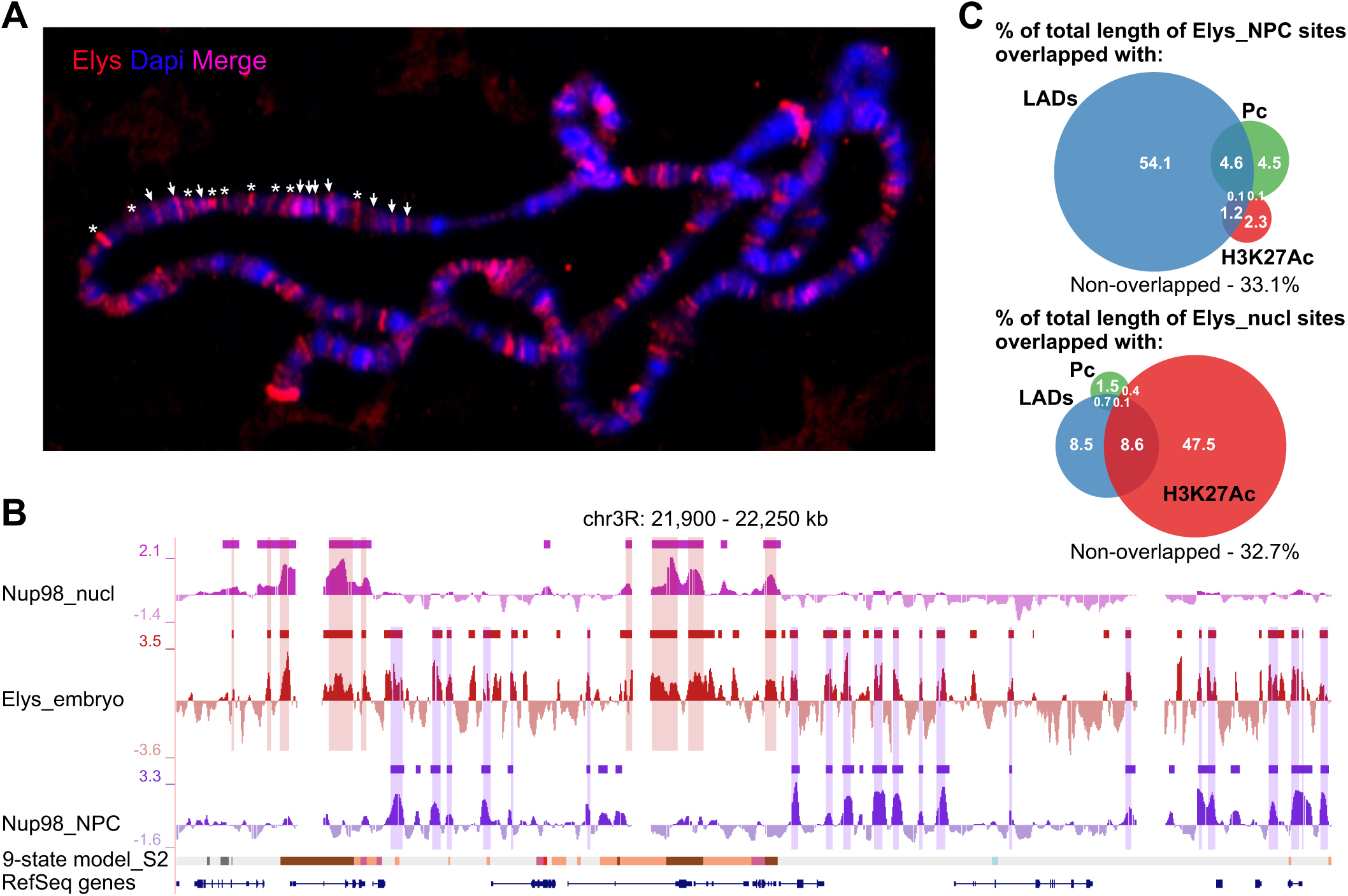
Elys interacts with chromatin located both at the NE and in the nuclear interior. (**A**) Immunostaining of squashed polytene chromosomes with anti-Elys (red) antibodies. Elys binding sites located within polytene chromosome bands, counterstained with Dapi (blue), are revealed as pink strips (marked by arrows in a chromosomal region), while Elys binding sites within inter-bands are revealed as red strips (marked by asterisks in a chromosomal region). (**B**) Screenshot from UCSC genome browser showing Nup98_nucl (pink), Elys_embryo (red) and Nup98_NPC (violet) profiles, as well as the corresponding domains or sites of enrichment (rectangles over profiles) for the representative region of chromosome 3R. The 9-state chromatin model and RefSeq genes are indicated below. The overlapped regions between Elys_embryo and Nup98_nucl sites or between Elys_embryo and Nup98_NPC sites are outlined by translucent rectangles. (**C**) Venn diagram showing the degree of overlap (as a percentage of Elys_NPC or Elys_nucl site length) between Elys_NPC or Elys_nucl sites and LADs, Pc and H3K27Ac domains.

### Identification of Elys binding sites in late embryos

We aimed to separate genomic sites bound by Elys in the nuclear interior from those bound by NPC-linked Elys. To this end, we employed publicly available data obtained by DamID technique [71] for Nup98-interacting regions in embryonic Kc167 cells which were successfully divided into two distinct pools, either interacting with Nup98 in the nucleoplasm (Nup98_nucl sites), or interacting with Nup98 at the NPCs (Nup98_NPC sites) [28]. Since almost all Nup98 ChIP-seq sites coincide with Elys ChIP-seq sites [35], we compared Elys ChIP-seq sites in S2 cells [37] with Nup98_nucl or Nup98_NPC sites in Kc167 cells (note that S2 and Kc167 are cognate cell cultures of embryonic origin). Surprisingly, while the overlap of Elys ChIP-seq sites with Nup98_nucl sites was highly non-random (*P* < 10^-4^, permutation test), the overlap of Elys ChIP-seq sites with Nup98_NPC sites was small and statistically non-significant (the overlapped regions constitute only 5.4% of total Elys peak coverage in the euchromatic chromosome arms, *P* ∼ 1 for their occasional colocalization, permutation test). Moreover, the overlap of Nup98 or Nup93 ChIP-seq sites identified in S2 cells [35,37] with Nup98_NPC sites determined in Kc167 cells was also non-significant (*P* ∼ 1, or *P* = 0.2, respectively, permutation test). The absence of colocalization between binding sites of different Nups identified by ChIP-seq in S2 cells with Nup98_NPC sites determined by DamID in Kc167 may be caused either by high variability of NPC-linked sites in various cell types, or by poor detection of NPC-bound chromatin by ChIP procedure.

To examine which assumption is correct, we tried to identify Elys binding sites in S2 cells using transient transfection of these cells with DamID constructs [72], but, by unknown reason, this approach gave unreliable results. Then, by applying DamID-seq technique adapted for transgenic flies [73], we successfully generated Elys profile in late (16-18-hour) embryos (Fig 2B). Using a three-state Hidden Markov model (HMM) algorithm, we identified ∼10600 Elys binding sites with median length 1.8 kb (S2 Table). Since late embryos represent a mixed population of diverse cell types, we reasoned that a fraction of Elys binding sites identified in late embryos might coincide with Nup98_NPC and Nup98_nucl sites in Kc167 cells. Indeed, we found that the majority of Elys peaks in embryos were colocalized either with Nup98_NPC or with Nup98_nucl peaks in Kc167 (Fig 2B). Whole-genome analysis revealed that ∼30% and ∼40% out of all Elys binding sites in embryos were highly non-randomly overlapped with Nup98_nucl and Nup98_NPC sites, respectively (*P* < 10^-4^ in both cases, permutation test). Hereinafter, these Elys binding sites conservative between embryos and Kc167 cells will be referred to as Elys_nucl and Elys_NPC sites (S2 Table). We also identified a fraction of ambivalent Elys_NPC/nucl sites (constituting ∼13% out of all Elys sites, S2 Table), apparently corresponding to Elys interactions with genome in both locations. Yet, ∼ 17% of Elys sites did not span Nup98 sites and were excluded from further analysis.

### NPC-linked Elys binds numerous genomic sites located within LADs

Recently, Gozalo *et al*. [37] have reported that 17% of Nup93 ChIP-seq sites and 22% of the shared Elys/Nup107 ChIP-seq sites in S2 cells were localized within LADs, thereby indicating peripheral positioning of a fraction of Nup-bound regions. Moreover, in some cases, positions of Nup93 ChIP-seq peaks coincided with the dips in the Lamin Dm0 (Lam) profile [37]. We determined the distribution of Elys_NPC and Elys_nucl sites across different types of chromatin. More than a half of Elys_NPC sites appear to locate within LADs (Fig 2C). In agreement with the reported overlap of some Nup93 binding sites with Pc domains [37], 9% of Elys_NPC sites were localized within Pc domains (Fig 2C). However, only a few Elys_NPC sites contain active, highly acetylated chromatin (Fig 2C). Strikingly, the degree of intersection (from the total length) of Elys_NPC sites with the inactive chromatin revealed in Kc167 cells [72] reaches 94%, while it was ∼70% with that revealed in S2 cells [74] (S4 Fig). Contrary to Elys_NPC sites, about a half of Elys_nucl sites overlap with H3K27 acetylated regions in S2 cells (Fig 2C), and 86% or 95% of Elys_nucl sites correspond to active chromatin types in Kc167 or S2 cells, respectively (S4 Fig). Therefore, Elys_NPC and Elys_nucl sites are almost completely located either within inactive or within active chromatin, respectively.

To validate that many Elys_NPC sites are indeed present within LADs, we compared Elys DamID profile in late embryos with Lam DamID profile in Kc167 cells [5]. In addition, we performed DamID-seq mapping of genomic regions interacting with Lam in 16-18-hour embryos, and identified embryonic LADs (S3 Table) covering 55.8% of non-repetitive genome.

Visual examination of the profiles in UCSC genome browser indicates that Elys_NPC peaks, positioned within LADs or at LAD boundaries, frequently correspond to the dips in the Lam profile from both Kc167 cells and late embryos (Fig 3A). Strikingly, the corresponding peaks were mostly absent in Elys ChIP-seq profile from S2 cells (Fig 3A). At the whole-genome level, the peak in the averaged Elys profile, centered at Elys_NPC sites positioned within LADs (±2 kb from LAD boundaries), corresponds to the dip in the averaged Lam profile from both Kc167 cells and late embryos (although in the latter case the dip is not so perfect and pronounced; Fig 3B). The dip is not revealed when the Lam profile is averaged around randomly chosen positions within Kc167 LADs (S5A Fig, left panel). Importantly, the dip in the Lam profile around Elys_NPC sites within LADs corresponds to the dip in the H3K27 acetylation profile (Fig 3B, most right panel). Therefore, Elys_NPC sites within LADs fail to interact with the NL not because of the increased histone acetylation correlating with chromatin positioning in the nuclear interior [70]. Rather, these genomic regions are bound by the NPC-linked Elys, and this binding partially prevents their interactions with the NL. In this case, the presence of Elys_NPC sites within LADs reflects the inaccuracy of two-state HMM algorithm, applied for LADs calling, which merges the neighboring LADs separated by short gaps corresponding to Elys_NPC sites.

**Fig 3.**
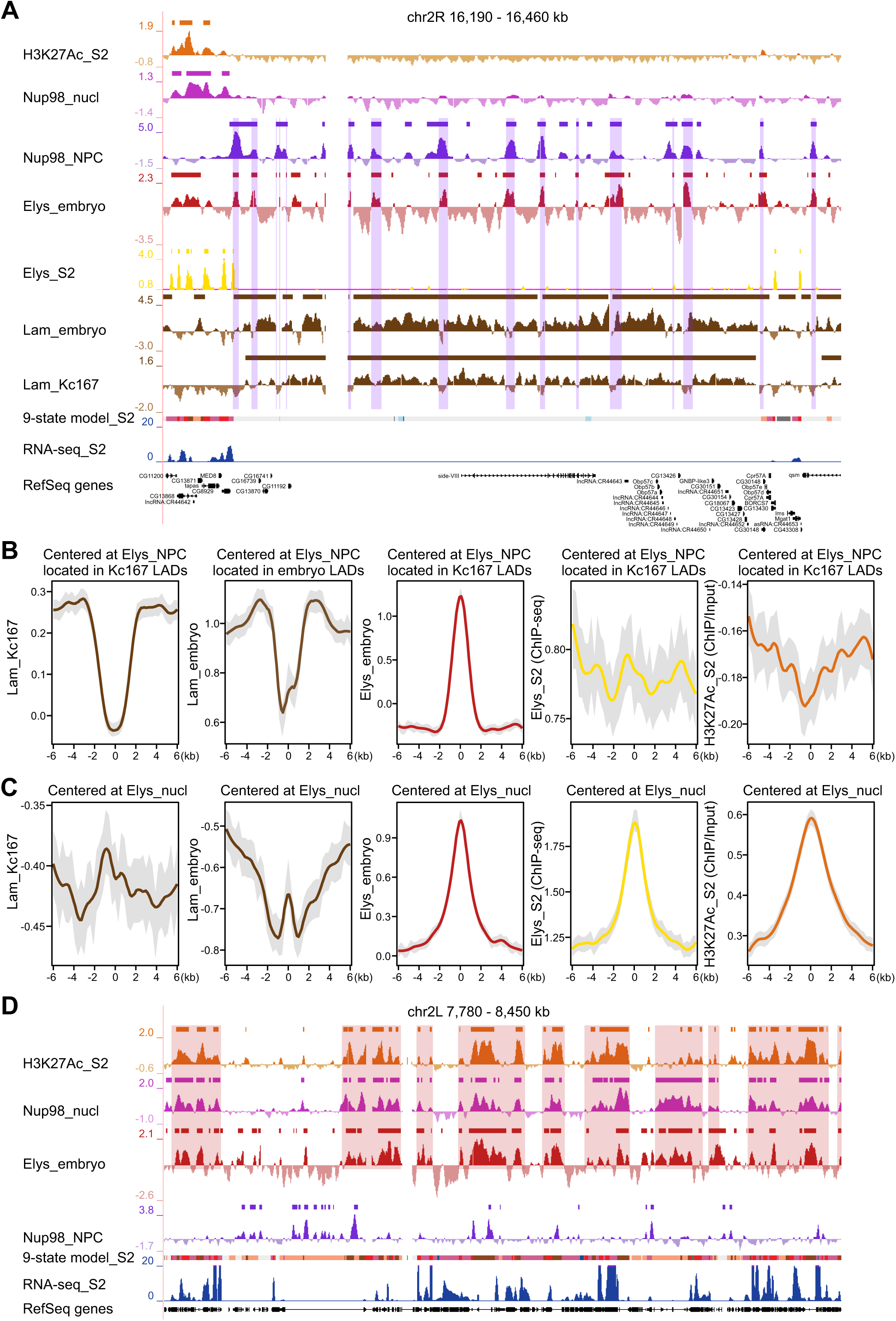
Elys_NPC sites are mostly located within LADs. (**A**) Screenshot from UCSC genome browser showing H3K27Ac (orange), Nup98_nucl (pink), Nup98_NPC (violet), Elys_embryo (red), Elys_S2 (yellow), Lam_embryo and Lam_Kc167 (brown) profiles, as well as the corresponding domains or sites of enrichment (rectangles over profiles) for the representative region of chromosome 2R. The 9-state chromatin model, RNA-seq in control S2 cells (from this work) and RefSeq genes are indicated below. The overlapped regions between Elys_embryo and Nup98_NPC sites containing Elys_NPC peaks are outlined by translucent rectangles (they mostly coincide with the dips in Lam profiles). (**B**,**C**) Averaged Lam_Kc167, Lam_embryo, Elys_embryo, Elys_S2 and H3K27Ac profiles around Elys_NPC sites (**B**) or Elys_nucl sites (**C**) located within LADs (±2 kb from LAD boundaries) determined in Kc167 or in late embryos. (**D**) Screenshot from UCSC genome browser showing H3K27Ac (orange), Nup98_nucl (pink), Elys_embryo (red) and Nup98_NPC (violet) profiles, as well as the corresponding domains or sites of enrichment (rectangles over profiles) for the representative region of chromosome 2L. The 9-state chromatin model, RNA-seq in control S2 cells (from this work) and RefSeq genes are indicated below. The regions of high concordance between H3K27Ac, Nup98_nucl and Elys_embryo profiles are outlined by translucent rectangles.

To explore whether the dips in the Lam profile are located at the same positions during development, we analyzed previously obtained Lam DamID profiles in the central brain, neurons, glia and the fat body of *Drosophila* third instar larvae [6]. The dips in the averaged Lam profile around Elys_NPC sites located within LADs (±2 kb from LAD boundaries) were detected in all these organs/cell types, including terminally differentiated neurons (S6 Fig). Therefore, during cell differentiation, NPC-linked Elys is bound to a similar set of genomic sites within LADs, although some variability associated with gene activation may be present.

It is noteworthy that the averaged Elys ChIP-seq profile from S2 cells [37] does not display the peak at Elys_NPC sites positioned within LADs (±2 kb from their boundaries) (Fig 3B). Moreover, the dip in the averaged Lam profile, centered at Elys ChIP-seq sites located within LADs (±2 kb from their boundaries) (S5B Fig), or within LADs overlapping with inactive chromatin from S2 cells (S5A Fig, right panel), is far less pronounced than that centered at Elys_NPC sites (S5A Fig, middle panel). Furthermore, in contrast to Elys_NPC sites, Elys ChIP-seq sites located within LADs (±2 kb from LAD boundaries) possess high level of histone acetylation (S5A Fig, most right panel). An increased histone acetylation is also revealed around Nup93 ChIP-seq sites from S2 cells (S5C Fig). The latter findings imply that the majority of these sites is likely located distantly from the NE in the active chromatin of inter-LADs adjoining to LAD boundaries, or in the transition zones between LADs and inter-LADs, but not within the genuine LADs.

Supporting this notion, Elys_nucl sites positioned in the nuclear interior demonstrate negative values of the averaged Lam profile and high level of histone acetylation (Fig 3C). High consistency between Elys_nucl profile and H3K27 acetylation is exemplified on Fig 3D.

Thus, we suppose that ChIP procedure may poorly reveal NPC-linked sites, whereas DamID technique readily detects them. The robust identification of Elys_NPC sites by DamID allows us to make a confident conclusion that peripheral chromatin interacts with the NE through multiple alternating sites. Some of them are attached to the NL, while others are attached to NPCs.

### Both NPC-linked and nucleoplasmic Elys likely recognize A/T-rich DNA motifs

Since Elys is the Nup known to bind chromatin directly [47,49,53–56], we analyzed DNA sequence motifs potentially responsible for its binding. Previously, Rasala *et al*. [47] have shown that antibiotic distamycin A, capable of binding with A/T-rich DNA sequences, inhibits *Xenopus* Elys interactions with chromatin. Furthermore, using EMSA, Mehta *et al*. [49] have revealed that two out of three non-canonical motifs in *Drosophila* Elys can bind A/T-rich sequences. Consistent with these results, we found that the most high-scoring motif identified by MEME [75] in Elys_NPC or Elys_nucl sites is represented by poly(A) tracks (Fig 4A). ∼30% of Elys_NPC and Elys_nucl sites contain at least one 20-bp A/T-rich sequence within ± 150 bp region from their centers. We also found that A/T-base content is increased in both Elys_NPC and, more notably, Elys_nucl sites compared to random genomic sites (Fig 4B). Therefore, a significant fraction of genomic targets of both NPC-linked and nucleoplasmic Elys contains A/T-rich DNA motifs, which likely mediate Elys binding to these sites.

**Fig 4.**
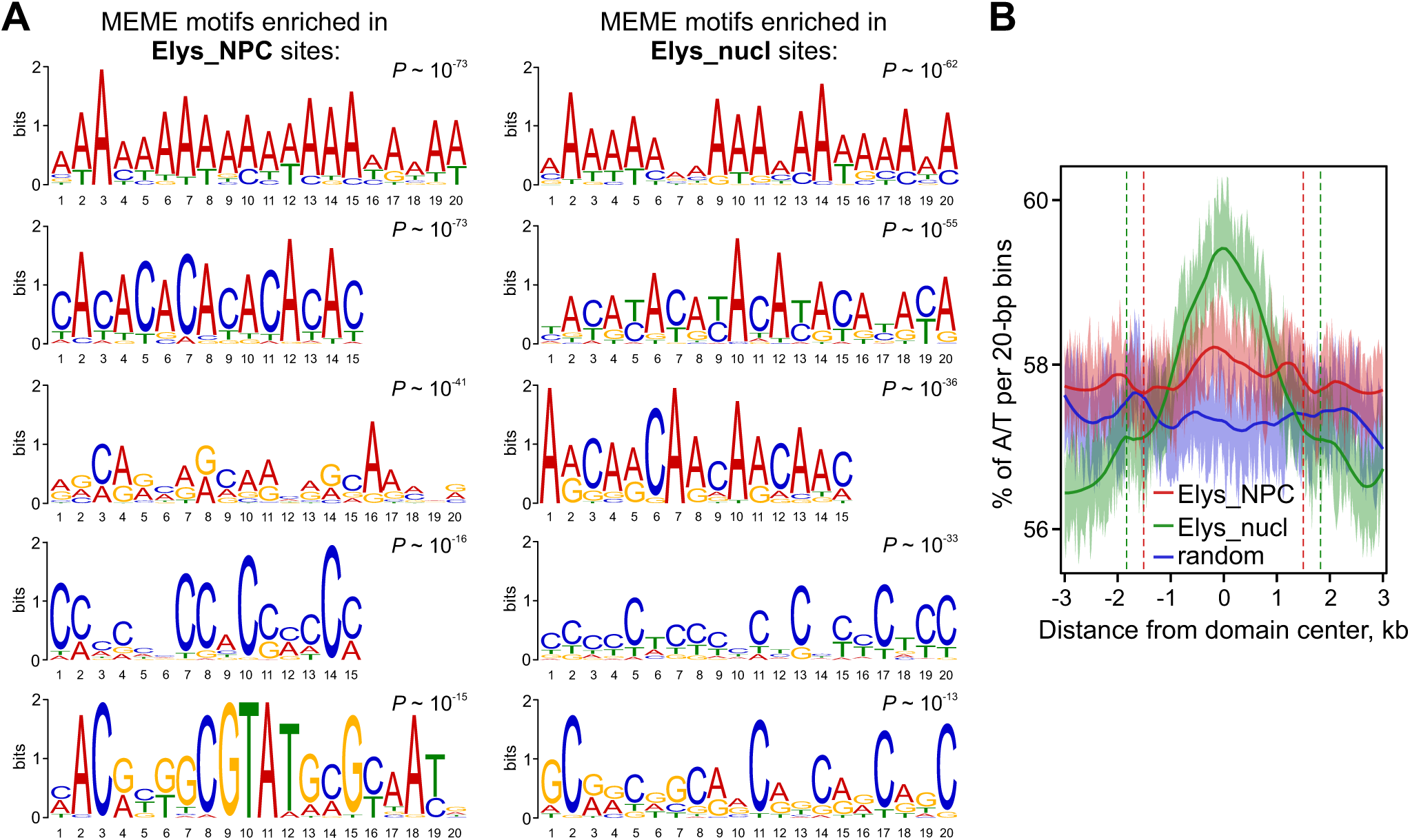
Elys sites possess an increased A/T content. (**A**) Top five MEME motifs identified in Elys_NPC or Elys_nucl sites (within ±150 bp from their centers). (**B**) Average A/T profile (percentage of A/T within 20-bp bins) around Elys_NPC, Elys_nucl or random genomic sites. Dashed lines delimit median lengths of corresponding sites.

### Binding of inactive chromatin with NPC-linked Elys maintains peripheral chromatin positioning at the NE

DamID technique detects genomic sites, which are either bound by or in contact with the protein of interest [76]. To confirm that Elys, as a component of NPCs, binds peripheral chromatin, but not just contacts it, we assayed by FISH the radial position of three loci, carrying Elys_NPC sites within LADs (S7 Fig), after Elys-KD in S2 cells (S1A and S1B Fig). These loci were previously shown to localize mostly at the NE and to lose peripheral localization upon Lam-KD in S2 cells [9,19]. We found that the lack of Elys leads to a notable displacement of these loci from the NE to the nuclear interior (Fig 5A–D, S4 Table), and this shift resembles that observed upon Lam-KD (Fig 5B). Moreover, simultaneous depletion of Elys and Lam causes a more noticeable removal of the locus from the NE (Fig 5B).

**Fig 5.**
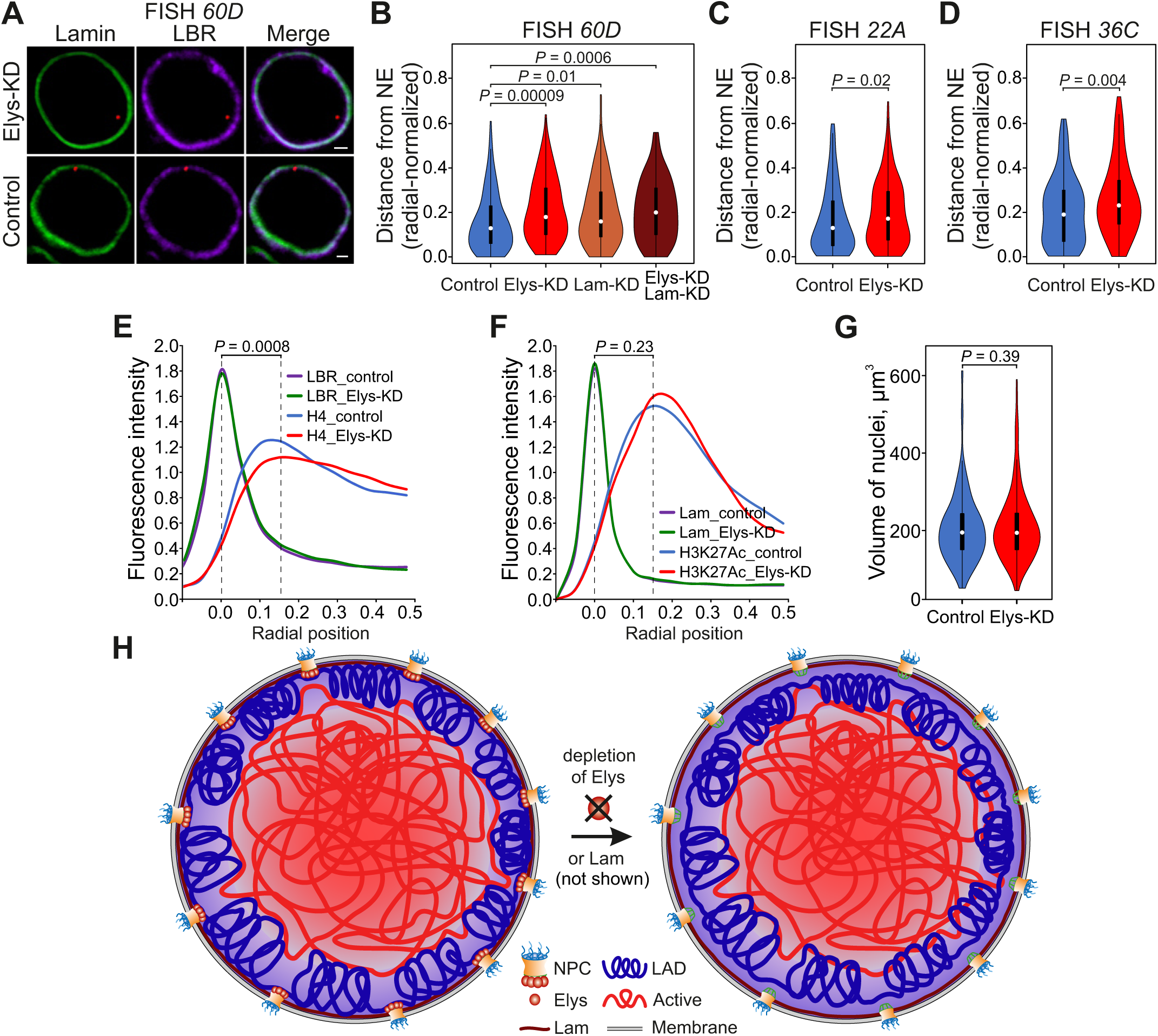
Chromatin is detached from the NE upon Elys-KD. (**A**) Confocal images of FISH signals (red) detected by the probe for *60D* region in nuclei stained with anti-Lam (green) and anti-LBR (violet) antibodies in Elys-KD or control S2 cells. Scale bar 1 µm. (**B**–**D**) Violin-plots showing distribution of radial-normalized distances between *60D* (**B**), *22A* (**C**), or *36C* (**D**) FISH signals and the NE in control (blue; n = 141, or n = 151, or n = 125 for *60D*, *22A* and *36C* probes, respectively), Elys-KD (red; n = 137, or n = 150, or n = 125 for *60D*, *22A* and *36C* probes, respectively), Lam-KD (brown; n = 100), and simultaneous Elys-KD and Lam-KD (dark brown; n = 100) S2 cells. (**E**,**F**) Averaged fluorescence intensity profiles along the nuclear diameter in Elys-KD (n = 50) and control (n = 50) S2 cells immunostained with anti-histone H4 and anti-LBR antibodies (**E**), or in Elys-KD (n = 170) and control (n = 170) S2 cells immunostained with anti-H3K27Ac and anti-Lam antibodies (**F**). (**G**) Violin-plots showing distribution of volumes of control (n = 250) and Elys-KD (n = 250) S2 nuclei manually outlined by the NL and further automatically reconstructed in *IMARIS*. (**H**) Scheme illustrating mechanisms of peripheral chromatin attachment to the NE. *P* values in (**B**-**G**) were estimated in a M-W *U*-test.

Next, we examined whether the removal of peripheral chromatin from the NE upon Elys-KD is a general phenomenon, or it affects only particular loci. To this end, methanol-fixed Elys-KD and control S2 cells were stained with anti-histone H4 antibodies to visualize total chromatin and with anti-LBR antibodies to visualize the NE. The averaged fluorescence intensity of histone H4, measured along the diameter of nuclei and normalized on the diameter of those nuclei, shows that total chromatin is localized slightly more distantly from the NE upon Elys-KD, as compared to control cells (Fig 5E, S8A Fig, S5 Table). However, staining of nuclei with anti-H3K27Ac antibodies indicates that active chromatin, located further away from the NE (S8B Fig), occupies nearly the same radial position in control and Elys-KD cells (Fig 5F, S5 Table). Therefore, the lack of Elys results in the inactive peripheral chromatin redistribution towards the nuclear interior.

Interestingly, volumes of nuclei were not significantly diminished upon Elys-KD compared to control S2 cells (Fig 5G, S6 Table), similar to what was observed upon Lam-KD in S2 cells [19].

Collectively, our findings indicate that, in interphase nuclei, inactive peripheral chromatin is attached not only to the NL, but also, via Elys, to the NPCs. Without each type of multiple anchorages, chromatin is slightly shifted from the nuclear periphery to the nuclear interior (Fig 5H).

### Elys-KD in S2 cells results in decompactization of the NE-attached TADs

Previously, the conflicting data were obtained regarding Elys influence on chromatin compaction. Artificial targeting of Elys via Sec13 to a particular site on polytene chromosomes was found to cause local chromatin decondensation [36]. However, ATAC-seq analysis upon Elys depletion in S2 cells did not reveal notable changes in chromatin accessibility at Elys ChIP-seq peaks or at all ATAC peaks [60]. To resolve this issue, we generated high-throughput chromosome conformation capture (Hi-C) [77] heatmaps at 4-kb resolution in control and Elys-KD S2 cells (S1C Fig). Using *Armatus* software [78], ∼3600 TADs in control and ∼3500 TADs in Elys-KD cells with 24-kb median length were identified (Fig 6A, S7 Table). Next, we calculated average contact frequency (ACF, see Materials and Methods) for each TAD that has exactly the same positions of boundaries in control and Elys-KD cells (∼57% of all control TADs). To explore TAD density changes upon Elys-KD, TADs with the same boundaries were divided into three groups depending on the proportion of coverage with both active chromatin (states 1 and 2 according to 9-state model in S2 cells [74]) and LADs [5]. For each TAD, the Jaccard coefficient based on these two metrics was calculated. After Elys-KD, TADs from group A, mostly composed of active chromatin, became slightly more compact, whereas TADs mostly corresponding to LADs (group C), on the contrary, became less densely organized (Fig 6B). The same trends in chromatin compaction were detected when TADs were divided into four groups based on quartiles of total gene expression within them (S9A Fig). Since the X chromosome in S2 cells is enriched in H4K16 acetylated histones due to the dosage compensation [79], we analyzed alterations in TAD density for the X chromosome separately and revealed the same ACF changes as for all TADs (S9B Fig). We conclude that the loss of Elys leads to more compact active TADs and less compact inactive TADs.

**Fig 6.**
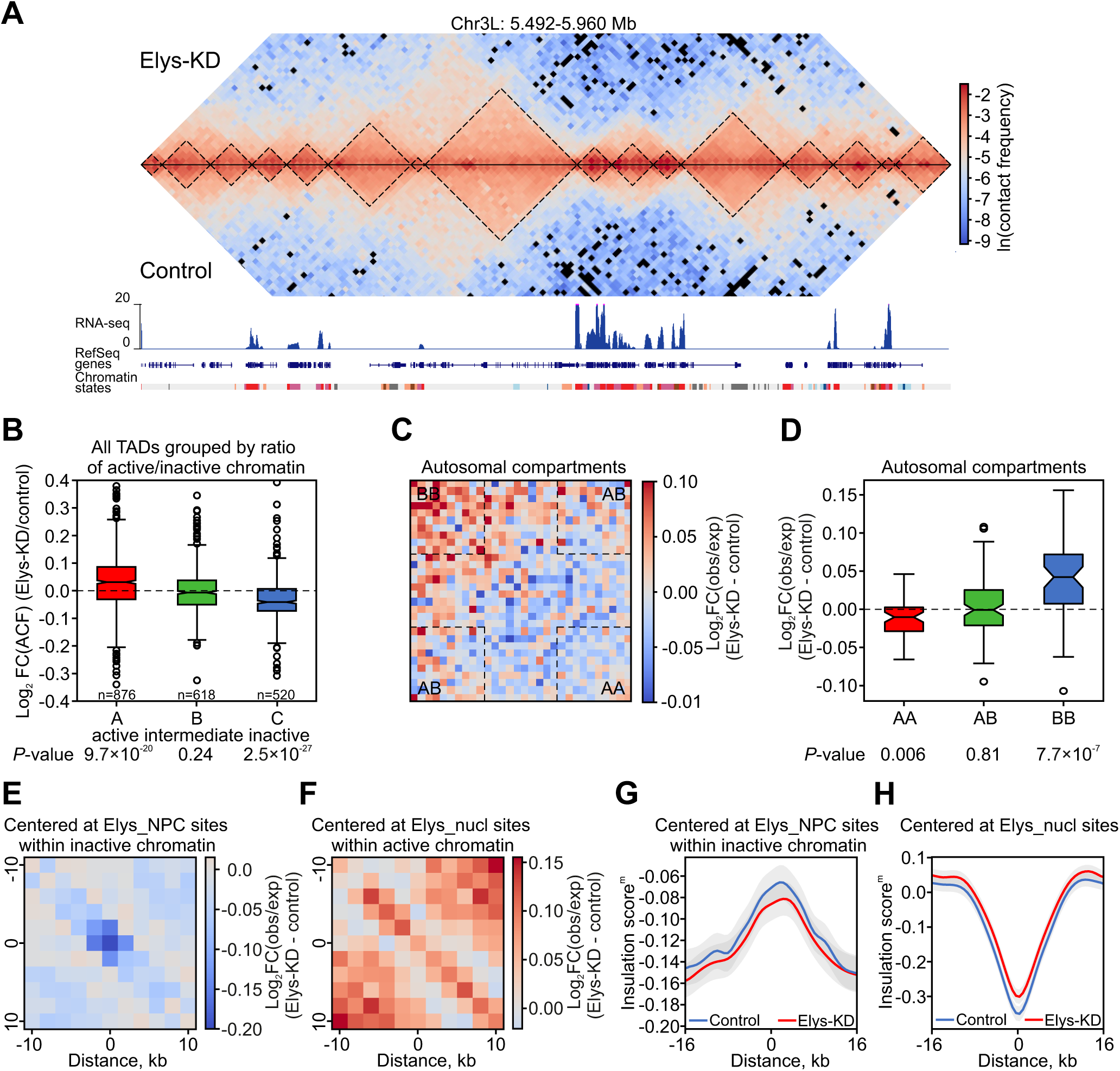
Hi-C analysis demonstrates more compact state of active chromatin and decompactization of inactive chromatin upon Elys-KD. (**A**) Hi-C heatmap for the representative region of 3L chromosome in Elys-KD (above the diagonal) and control (below the diagonal) cells. TADs (demarcated by black dotted lines), RNA-seq profile in control cells (in RPM, blue peaks), RefSeq genes, and chromatin annotation in S2 cells according to the 9-state model are indicated below the Hi-C map. (**B**) Log_2_ FC of ACF (Elys-KD/control) for TADs divided into three groups (A, B, C). *P* values were estimated in a Wilcoxon signed-rank test. (**C**) Saddle plot showing subtraction of aggregated contact frequency (Elys-KD minus control) log_2_FC(observed/expected) for autosomes. Dashed lines indicate areas which were analysed on the panel (**D**). (**D**) Box-plots showing subtraction of aggregated contact frequency (Elys-KD minus control) log_2_FC(observed/expected) for autosomes within active (AA), inactive (BB), and between active and inactive (AB) chromatin compartments. *P* values were estimated in a Wilcoxon signed-rank test. (**E**,**F**) Heatmaps showing subtraction of ACF (Elys-KD minus control) log_2_FC(observed/expected) around Elys_NPC sites located in the inactive chromatin (states 6-9) (**E**) or around Elys_nucl sites containing > 50% of active chromatin (states 1 and 2) (**F**). (**G**,**H**) Averaged IS^m^ profiles around Elys_NPC sites located within inactive (states 6-9) chromatin (**G**) or around Elys_nucl sites (**H**) in control (blue curve) and Elys-KD (red curve) cells. IS^m^ profiles were calculated for 2-kb resolution Hi-C heatmaps. Only contacts between regions located 2-8 kb away from both sides of the central bin were considered.

Next, we analyzed distant interactions within and between active (A) and inactive (B) chromatin compartments [77]. As was reported earlier [19], contact frequency in S2 cells is higher than expected only for the A compartment (S9C Fig). Similar to the impact of Lam-KD [19], Elys-KD results in enhanced or relaxed distant interactions within inactive or active autosomal compartments, respectively (Fig 6C and 6D). The same trend is observed for the inactive compartment of the X chromosome (S9D Fig). Yet, the enhanced intermingling of A and B compartments reported upon Lam-KD, is not observed upon Elys-KD.

To track the influence of Elys on local chromatin compaction, we built averaged Hi-C heatmaps around Elys_NPC and Elys_nucl sites. Upon Elys-KD, ACF within Elys_NPC sites in inactive chromatin (states 6-9 [74]) decreases stronger than within adjacent inactive regions (*P* = 0.002, M-W U-test) (Fig 6E). Moreover, chromatin attachment to NPCs facilitates contacts between regions located 2-8 kb away from both ends of Elys_NPC sites, as determined by the peak on the curve of modified insulation score (IS^m^, see Materials and Methods for details) reflecting contact frequency between these regions (Fig 6G). Such contacts become weaker upon Elys-KD (*P* = 9.1×10^-7^, Wilcoxon signed-rank test). These results indicate a peculiar three-dimensional organization of Elys_NPC sites, which is likely mediated by their attachment to NPCs.

At the same time, ACF within Elys_nucl sites containing ≥ 50% of active chromatin (states 1 and 2 [74]) as well as within adjacent active regions is increased upon Elys-KD (Fig 6F). Elys occupancy of Elys_nucl sites, which coincide with a fraction of TAD boundaries [80], enhances spatial isolation of neighboring regions (*P* = 1.5×10^-61^, Wilcoxon signed-rank test) (Fig 6H). Therefore, Elys binding with active chromatin leads to its slight decompactization, which may be mediated by the PBAP complex, shown to be associated with Elys [36].

### Elys is enriched at the 5’- and 3’-ends of genes

Nups in mammals and *Drosophila* were found to interact with enhancers or super-enhancers (the latter represent enhancer clusters) [81], marked by increased H3K27 acetylation [34,35]. Using modified DamID technique, super-enhancers in mammals were recently shown to associate with the NPC-linked Nups [82]. It remained unclear whether enhancers in *Drosophila* are also associated with NPCs. We examined whether Elys_nucl or Elys_NPC sites are colocalized with strong active enhancers which were identified in S2 cells by self-transcribing active regulatory region sequencing (STARR-seq) [83]. We found highly statistically significant overlap only between STARR-seq enhancers and Elys_nucl sites (Fig 7A; 42% of enhancers overlap with Elys_nucl sites, *P* < 10^-4^ for their occasional colocalization, permutation test), but not between STARR-seq enhancers and Elys_NPC sites (5% of enhancers overlap with Elys_NPC sites, *P* ∼ 1 for their occasional colocalization, permutation test). However, there was still a possibility that enhancers induced by external stimuli may be attached to the NPCs. To test this idea, we compared Elys_nucl and Elys_NPC sites with STARR-seq enhancers activated by ecdysone treatment in S2 cells [84]. Similar to active enhancers, the ecdysone-responsive enhancers strongly colocalize with Elys_nucl sites, but not with Elys_NPC sites (Fig 7A; *P* < 10^-4^ and *P* ∼ 1, respectively, for their occasional colocalization, permutation test). Therefore, both active and inducible enhancers in *Drosophila* S2 cells are mostly bound by the nucleoplasmic Elys fraction.

**Fig 7.**
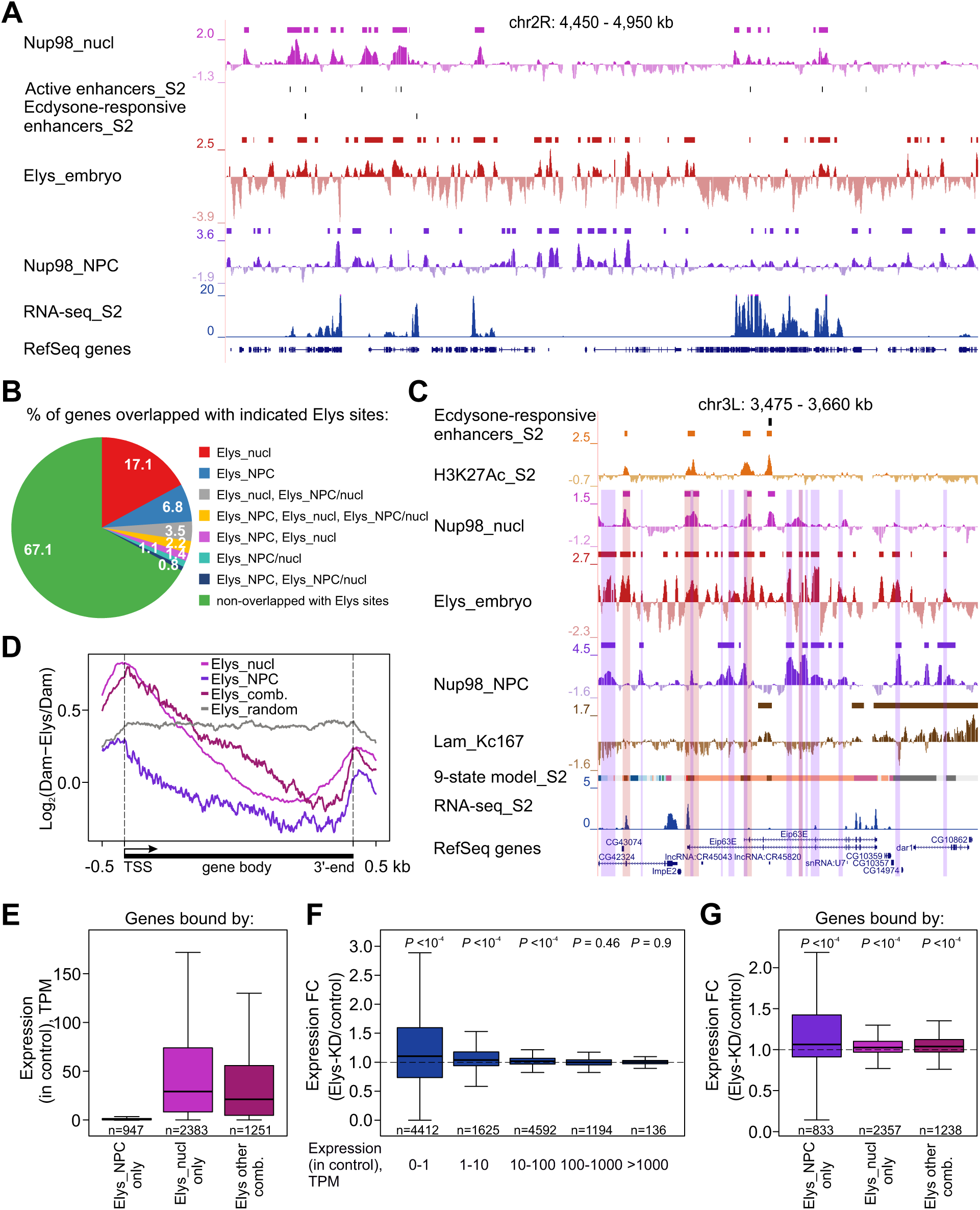
Elys weakly affects gene expression. (**A**) Screenshot from UCSC genome browser demonstrating that active or ecdysone-responsive STARR-seq enhancers mainly coincide with Elys_nucl sites. Nup98_nucl (pink), Elys_embryo (red), and Nup98_NPC (violet) profiles are shown, as well as the corresponding sites of enrichment (rectangles over profiles) and active or ecdysone-responsive STARR-seq enhancers in S2 cells (black rectangles) for the representative region of chromosome 2R. RNA-seq in control S2 cells (this work) and RefSeq genes are indicated below. (**B**) Pie chart showing the percentage of genes overlapping with indicated Elys sites. (**C**) Screenshot from UCSC genome browser with an example of gene containing Elys_nucl, Elys_NPC and Elys_NPC/nucl sites. H3K27Ac (orange), Nup98_nucl (pink), Nup98_NPC (violet), Elys_embryo (red) and Lam_Kc167 profiles are shown, as well as the corresponding domains or sites of enrichment (rectangles over profiles) and ecdysone-responsive STARR-seq enhancers in S2 cells (black rectangles). RNA-seq in control S2 cells (this work) and RefSeq genes are indicated below. Regions corresponding to Elys_NPC, Elys_nucl, and Elys_NPC/nucl sites are outlined by violet, pink or merged color translucent rectangles, respectively. (**D**) Averaged Elys profiles over metagene carrying sites of Elys_NPC (violet), Elys_nucl (pink), their combinations (wine-colored), or for randomly reshuffled genes (grey). (**E**) Box-plots showing RNA-seq expression (in TPM) in control S2 cells for genes carrying sites of Elys_NPC (violet), Elys_nucl (pink), or their combinations (wine-colored). (**F**) Box-plots showing RNA-seq expression FC (Elys-KD/control) ranked by the expression level in control S2 cells. (**G**) Box-plots showing RNA-seq expression FC (Elys-KD/control) for genes carrying sites of Elys_NPC (violet), Elys_nucl (pink), or their combinations (wine-colored). *P* values in (**F**,**G**) were estimated in a Wilcoxon signed-rank test.

17.1% and 6.8% of genes overlap solely with Elys_nucl or Elys_NPC sites (Fig 7B, S8 Table), thus indicating the prevalent positioning of these genes in the nuclear interior or at the NE, respectively. However, 8.2% of genes simultaneously contain various combinations of these sites (including Elys_NPC/nucl sites) (Fig 7B, S8 Table). The ecdysone-inducible gene *Eip63E* (Fig 7C) belongs to the latter group of genes. They are attached to NPCs by internal sites (which in some cases include enhancers) and are looped out from NPCs by other regions. Such an organization may be associated with the transcriptional memory, inherent to ecdysone-inducible genes [35].

Elys_NPC sites are randomly distributed between genes and intergenic regions, whereas Elys_nucl sites are predominantly localized at gene promoters (S10 Fig). The metagene profile demonstrates that Elys occupies mostly 5’- and 3’-ends of genes (Fig 7D). To examine possible mechanism(s) of this enrichment, we built averaged metagene profiles for A/T content, H3K27 acetylation and binding of GAF. The peak of Elys binding at the 3’-ends of genes exactly coincides with peaks of A/T content and H3K27 acetylation (S11A and S11B Fig). Thus, Elys enrichment at the 3’-ends of genes may be mediated by either of these features. However, Elys peak at TSSs coincides with the peak of GAF, but not with peaks of A/T content or H3K27 acetylation (S11C Fig). Since GAF may guide the PBAP complex to TSSs [59,60], Elys that was shown to associate with the PBAP [36], may be recruited to TSSs via this mechanism. Yet, we did not find that Elys and GAF are co-immunoprecipitated from S2 cells (S11D Fig).

### Elys-KD in S2 cells causes derepression of genes in LADs

To explore the influence of Elys on gene expression, we performed poly(A^+^) RNA-seq analysis in the control and Elys-KD S2 cells (S1D Fig, S8 Table). Unexpectedly, changes in gene expression upon Elys-KD were rather weak. We identified only 175 differentially expressed genes (i.e., genes with fold change (FC) of expression > 1.5 and false discovery rate < 0.05). 131 genes were up-regulated and 44 genes were down-regulated (S8 Table). Our further analysis was focused on genes bound by NPC-linked or nucleoplasmic Elys. Consistent with low and high levels of histone acetylation at Elys_NPC and Elys_nucl (or Elys_NPC/nucl) sites, respectively, genes carrying only Elys_NPC sites were mostly “silent”, while genes carrying only Elys_nucl sites or various Elys site combinations were actively expressed (Fig 7E). Elys-KD resulted in slightly increased expression of “silent” and weakly expressed genes, while most active genes did not notably change their expression (Fig 7F). Accordingly, genes carrying only Elys_NPC sites were slightly up-regulated, while genes carrying Elys_nucl sites or various Elys site combinations were barely affected upon Elys-KD (Fig 7G). We note that the majority of “silent” or weakly expressed genes, that were up-regulated upon Elys-KD and either contained or not contained Elys_NPC sites, were localized within LADs. Together with our previous results showing that the same effect on gene expression was detected upon Lam-KD [19], these data support the idea that up-regulation of weakly expressed genes upon Elys-KD may be caused by the loss of their interactions with the NL. We conclude that Elys-KD faintly affects gene expression.

## Discussion

### Elys-independent post-mitotic NPC incorporation in the NE

In this work, we found that *Drosophila* S2 cells is an appropriate model to analyze the influence of Elys on overall genome architecture, since, unlike in other organisms and some other *Drosophila* cell types [42–46,49], depletion of Elys in S2 cells does not result in a notable reorganization of NE components, including NPCs (Fig 1A–1C), and does not cause the enhanced apoptosis (S3 Fig). Moreover, upon Elys depletion in S2 cells, post-mitotic NPC incorporation in the NE occurs prior to telophase (Fig 1H). Then, what might be a mechanism of rapid post-mitotic NPC incorporation in the absence of Elys? There is an Elys-independent pathway for insertion of NPCs in the NE during interphase, but it is rather slow, at least in mammals [48]. However, in early *Drosophila* embryos, the pre-assembled NPCs, associated with membrane stacks of the endoplasmic reticulum (with the annulate lamellae) [85], are incorporated into the reforming NE shortly after mitosis [86]. Moreover, similar mechanism operates in various mammalian cell lines as well as in the differentiated *Drosophila* cells [87]. We hypothesize that, upon Elys-KD, embryonic S2 cells may restore NPCs at the NE at the end of mitosis by employing the same mechanism.

### Mechanisms of Elys binding with chromatin

Using DamID technique, we identified Elys binding sites in late embryos. By comparing them with genomic regions interacting with Nup98 either in the nucleoplasm or at the NPCs [28], we identified conservative Elys_NPC and Elys_nucl sites. Elys_NPC sites are largely preserved during cell differentiation, contain non-acetylated nucleosomes and are located within LADs or, less frequently, within Pc domains (Figs 2C, 3A and 3B). On the contrary, Elys_nucl sites contain acetylated chromatin (Figs 2C, 3C and 3D) and are colocalized with active gene promoters (S10 Fig) and enhancers. Interestingly, both Elys_NPC and Elys_nucl sites bear A/T-rich DNA motifs (Fig 4) likely determining their recognition by Elys. The prevalence of sequence-specific mode of Elys binding is supported by the fact that the majority (83%) of Elys binding sites in late embryos are shared with Nup98 binding sites in Kc167 cells.

We hypothesize that, in anaphase, Elys recognizes and binds the A/T-rich sequences within decondensing chromatin and maintains binding with them during interphase. In this scenario, whether a particular Elys molecule appears at the NE (within NPC) or in the nucleoplasm depends on whether its A/T-rich binding site is embedded in the non-acetylated or acetylated chromatin, respectively. In the acetylated chromatin, Elys binding with A/T-rich sequences may promote Elys spreading on the neighboring acetylated regions due to Elys ability to associate with the acidic patch of H2A/H2B histone dimer [55,56]. This ability may explain the reported correlation between Elys and H3K27 acetylation profiles [35] (Fig 3D). Yet, Elys association with the H2A/H2B acidic patch is likely to be blocked in non-acetylated nucleosomal arrays by the non-acetylated N-terminal tail of histone H4 competing for binding with the same interface [88]. An additional level of complexity in Elys interactions with chromatin is mediated by Elys targeting to some genomic sites by its protein partners. In particular, the colocalization of Elys and GAF peaks at gene promoters (S11C Fig) may suggest an existence of indirect mechanism of Elys targeting to chromatin via GAF/PBAP complex [36,58,59].

### Elys establishes and maintains peripheral chromatin positioning

We reason that if Elys links chromatin with NPCs upon mitotic exit and interacts with chromatin in interphase nuclei, it could be required to keep peripheral chromatin attached to the NE during interphase. Indeed, using FISH and chromatin immunostaining we found that several loci analyzed as well as peripheral chromatin as a whole were relocated from the NE to the nuclear interior upon Elys-KD in S2 cells (Fig 5). This effect resembles the one observed upon Lam-KD in S2 cells [19]. Our results indicate that NPC-linked Elys is not simply in contact with chromatin, but is bound to it. Moreover, NPC-linked Elys maintains localization of peripheral chromatin at the NE. Elys binding to peripheral chromatin is also supported by the dips in Lam profiles at Elys_NPC sites (Fig 3A and 3B), as well as by the peculiar three-dimensional organization of these sites (Fig 6E and 6G). Taken together, our findings suggest that peripheral chromatin is simultaneously bound both to the NL and, via Elys, to the NPCs (Fig 5H). As far as we know, our results provide the first evidence that peripheral chromatin in a multicellular organism is maintained at the NE not only due to its interactions with the NL, but also due to its interactions with NPCs. These findings refute a model postulating that chromatin associated with nuclear pores constitutes islets of active genes within the repressed chromatin layer lining the nuclear envelop.

### Impact of Elys on chromatin compaction and transcription

Apart from the maintenance of genome architecture, NPC-linked Elys binds silent, or weakly expressed genes, whereas nucleoplasmic Elys binds promoters and enhancers of actively expressed genes. Our RNA-seq analysis shows that Elys-KD in S2 cells only weakly affects gene expression. Low-expressed genes become mainly up-regulated, while transcription of active genes is mostly unaltered (Fig 7E–7G). The similarity of effects on gene expression upon Elys-KD and Lam-KD [19] implies that the derepression of low-expressed genes upon Elys-KD may be caused by the detachment of genes from the NL, but not by direct impact of Elys on their transcription. In support of this notion, we found that TADs attached to the NL become decompacted upon both Elys-KD (Fig 6B) and Lam-KD [19]. At the same time, active TADs become more compact after Elys-KD (Fig 6B), while transcription within them is mostly unchanged (Fig 7E–7G). Therefore, if Elys recruits the PBAP chromatin remodeling complex to the active genes [36], this provides only the potential for them to be activated. Of note, Elys binding with promoters and enhancers may be related to the transcriptional memory, i.e., more rapid and strong induction of gene transcription upon repeated treatment of cells with external activation stimuli. In metazoans, this phenomenon was associated with several Nups including Nup98 and Nup153 [35,89,90].

In conclusion, our findings point to the key role that Elys, as a constituent of NPCs, plays in the maintenance of peripheral positioning of inactive chromatin during interphase. Elys is also associated with the promoters and enhancers of active genes in the nuclear interior, resulting in their chromatin opening. However, it does not notably affect gene expression, and its functional significance remains to be explored.

## Materials and methods

### Plasmid construction

To generate *pUAST-attB-hsp70-loxP-stop-loxP-Dam-Elys* construct, the whole *Elys* ORF, lacking start codon, was initially assembled in the *pBluescript II* vector (Stratagene). To this end, the ∼0.32-kb 5’-fragment and the ∼1.0-kb 3’-fragment of *Elys* ORF were amplified by PCR from cDNA LD14710 (Drosophila Genomics Resource Center) using primers indicated in S9 Table. Then, NotI- and NheI-digested 5’-fragment, HindIII- and ApaI-digested 3’-fragment and ∼5.1-kb NheI- and HindIII-digested central fragment of *Elys* ORF were combined together by cloning into the HindIII and ApaI sites of *pBluescript II*. Next, the whole *Elys* ORF was excised by NotI and SpeI sites and cloned by NotI and XbaI sites in *pUAST-attB-hsp70-loxP-stop-loxP-Dam-Lam* construct [91] instead of Lamin *Dm0* ORF giving *Elys* ORF in frame with *Dam* ORF. The correct nucleotide sequence of resultant *pUAST-attB-hsp70-loxP-stop-loxP-Dam-Elys* construct was verified by partial sequencing.

### Fly stocks and handling

Fly stocks were maintained under standard conditions at 25°C. Transgenic strains carrying *pUAST-attB-hsp70-loxP-stop-loxP-Dam-Elys* were generated by *ϕC31*-mediated site-specific integration at the *attP40* site on the chromosome 2 in the *y*, *w*; *P{y[+t7.7] = CaryP}attP40*; *M{3xP3-RFP.attP}ZH-86Fb*; *M{vas-int.B}ZH-102D* line [92] as previously described [93]. Transgenic strains carrying *pUAST-attB-hsp70-loxP-stop-loxP-Dam*, *pUAST-attB-hsp70-loxP-stop-loxP-Dam-Lam* and *pUAST-attB-nos-Cre* constructs at the same genomic site were described previously [65,73,91].

### DamID-seq procedure in late embryos

To perform DamID in late embryos, we crossed females expressing Cre-recombinase in the oocytes and zygotes (#766 line from the Bloomington Drosophila Stock Center which was devoid from balancers) with the males carrying *pUAST-attB-hsp70-loxP-stop-loxP-Dam* or *pUAST-attB-hsp70-loxP-stop-loxP-Dam-Lam* or *pUAST-attB-hsp70-loxP-stop-loxP-Elys* constructs. These constructs contain stop-cassette, flanked by *loxP* sites, which separates basal *hsp70* promoter from *Dam*, *Dam-Lam* or *Dam-Elys* ORFs [73]. An excision of stop-cassette in early embryos results in Dam-methylation of corresponding genomic sites in late embryos. Approximately two hundred 16-18-h embryos from these crosses were collected, washed with H_2_O, dechorionated with 2.5% hypochlorite for 3 min and again washed three times with H_2_O. Further isolation of genomic DNA, amplification of Dam-methylated genomic fragments and their subsequent sequencing were performed according to [94]. 16 (for *Dam-Lam*) or 18 (for *Dam* and *Dam-Elys*) cycles of PCR amplification (1 min at 94°C, 1 min at 65°C, 2 min at 68°C) was applied for DNA samples. Sequencing on Illumina HiSeq 2500 was performed in the Evrogen facility (www.evrogen.ru) resulting in 27–30 million 100-nt single-end reads per sample (S10 Table).

### Immunostaining of polytene chromosomes

Polytene chromosomes staining was performed according to [95] with some modifications. Chromosomes were stained with anti-Elys antibodies [65] (1:400) diluted in PBT containing 0.3% Triton X-100 and 3% Normal Goat Serum (Invitrogen) and counterstained with Dapi. As the secondary, Alexa Fluor 546-conjugated goat anti-rabbit IgG (1:200; Invitrogen) was used. Image capture was performed with a confocal *LSM 510 Meta* laser scanning microscope (Zeiss).

### Cell culture maintenance and RNAi

*Drosophila melanogaster* S2 cell line (from IMG collection) was grown at 25°C in Schneider’s Drosophila Medium (Gibco) supplemented with 10% heat-inactivated fetal bovine serum (FBS, Gibco), 50 units/ml penicillin, and 50 µg/ml streptomycin. For RNAi treatment of S2 cells, dsRNAs against *LacZ* (control), *Lam*, or *Elys* were prepared as was described previously [9] using primers indicated in S9 Table. Cells were treated with dsRNAs during four days, or during three days with an additional treatment for two days using previously described protocol [96].

### Western-blot analysis

Proteins were extracted with 8 M urea, 0.1 M Tris-HCl, pH 7.0, 1% SDS, fractionated by SDS-PAGE (12% acrylamide gel) and transferred to a PVDF membrane (Immobilon-P, Millipore). Blots were developed using alkaline phosphatase-conjugated secondary antibodies (Sigma) and the Immun-Star AP detection system (Bio-Rad). The following antibodies were used for detection: mouse monoclonal anti-Lam [97] (1:2000; ADL67), mouse monoclonal anti-beta Actin (1:5000; ab8224, Abcam), rabbit polyclonal anti-Nup107 [98] (1:5000), rabbit polyclonal anti-Elys [65] (1:3000).

### Immunostaining of S2 cells

S2 cells in the growth phase were collected and rinsed two times in PBS. Cells were fixed in 4% formaldehyde (in PBT) for 25 min at room temperature. Fixation was stopped by incubation with 0.25 M glycine (Sigma-Aldrich) for 5 min. Further immunostaining procedure was performed as previously described [6]. As the primary, mouse monoclonal anti-Lam [97] (ADL84, 1:500), guinea pig polyclonal anti-LBR [99] (1:1000), mouse monoclonal Mab414 (1:300; Abcam ab24609), chicken polyclonal anti-CenpA [100] (1:600), rabbit polyclonal α-Tubulin (1:2000; Abcam ab18251) and rabbit polyclonal anti-Elys [65] (1:1000) antibodies were used. As the secondary, Alexa Fluor 488-conjugated goat anti-rabbit IgG (Invitrogen), or Alexa Fluor 488-conjugated or 633-conjugated goat anti-mouse IgG (Invitrogen), or Alexa Fluor 633-conjugated goat anti-chicken IgG (Invitrogen) antibodies were used.

### TUNEL assay

TUNEL assay was performed using Click-iT Plus TUNEL kit (C10619, Invitrogen). After seeding on the coverslip, fixation in 4% formaldehyde (in PBT) for 25 min at room temperature, stopping the reaction with 0.25 M glycine (Sigma-Aldrich) for 5 min and performing Click-iT Plus reaction according to manufacturer’s instructions, control and Elys-KD S2 cells were immunostained with mouse monoclonal anti-Lam [97] (ADL84, 1:500) and rabbit polyclonal anti-Elys [65] (1:1000) antibodies. Confocal images were captured using *LSM 710* laser scanning microscope (Zeiss). The percentage of apoptotic cells in control or Elys-KD cells was counted for 10 images per replicate in three biological replicates (S1 Table).

### DNA FISH

Hybridization probes for the *22A* and *36C* regions were prepared using long-range PCR with primers indicated in S9 Table. Cosmid clone k9 [9] was used as hybridization probe for the *60D* region. FISH with S2 cells was performed as was described previously [19]. As the primary antibodies, we used guinea pig polyclonal anti-LBR [99] (1:1000), mouse monoclonal anti-Lam [97] (ADL84, 1:500), rabbit polyclonal anti-Elys [65] (1:1000), sheep polyclonal anti-DIG-FITC (1:500, Roche). As the secondary antibodies we used Alexa Fluor 633-conjugated goat anti-guinea pig IgG (Invitrogen), Alexa Fluor 488-conjugated or 633-conjugated goat anti-mouse IgG (Invitrogen), Alexa Fluor 488-conjugated goat anti-rabbit IgG (Invitrogen), Alexa Fluor 488-conjugated goat anti-FITC IgG (Invitrogen).

### RNA FISH with oligo(dT) probe

S2 cells were fixed with 3.7% formaldehyde in PBS for 10 min. Fixation was stopped by incubation with 0.25 M glycine (Sigma-Aldrich) for 5 min. Next, cells were washed in PBS, permeabilized for 10 min with PBS containing 0.5% Triton X-100 and incubated for 15 min at 37°C in prehybridization buffer (2× SSC, 20% formamide, 0.2% BSA, 1 mg/ml tRNA). Hybridization was performed overnight at 37°C in the prehybridization buffer supplemented with 1 µM oligo(dT)_40_- Cy5. The cells were washed twice for 5 min with 2× SSC, 20% formamide at 42°C, once with 2× SSC at 42°C, once with 1× SSC and once with PBS at room temperature. Next, cells were immunostained with mouse monoclonal anti-Lam [97] (ADL84, 1:500) and rabbit polyclonal anti-Elys [65] (1:1000) antibodies.

### Measuring distances from FISH signals to the NE

Three-dimensional image stacks were recorded with a confocal *LSM 510 Meta* or *LSM 710* laser scanning microscope (Zeiss). Optical sections at 0.35-0.4 μm intervals along the Z-axis were captured. Images were processed and analyzed using *IMARIS 7.4.2* software (Bitplane AG) with a blind experimental setup. Images were thresholded to eliminate non-specific background. The distances between signals and nuclear envelope were counted as previously described [9]. Briefly, NL stained by anti-LBR antibodies (for FISH with *60D* and *22A* probes) or by anti-Lam antibodies (for FISH with *22A* and *36C* probes) was manually outlined by its middle in each plane of the Z-stack, before automatic reconstruction of the nuclear surface and calculation of nuclear volume (S6 Table) by *IMARIS*. One measurement point was positioned in the optical section with the brightest FISH signal, at its visually determined center, and another one was placed on the reconstructed nuclear surface at the point of its earliest intersection with the progressively growing sphere from the first measurement point. The distance between the measurement points (the shortest distance between the center of FISH signal and the middle of the NL) was measured for each nucleus. Data were obtained in two or three replicates with 35-100 FISH signals per replicate (S4 Table). Distances were normalized on the nuclei radii, which were calculated from the volumes of reconstructed nuclei on the assumption that they have the spherical form.

### Chromatin visualization by histone H4 or H3K27Ac

Elys-KD or control S2 cells were seeded on coverslips for 30 min. After rinsing with PBS, cells were fixed in 100% methanol for 5 min at room temperature, rinsed with PBS three times and blocked with PBTX (PBS with 0.1% Tween-20 and 0.3% Triton X-100) containing 3% normal goat serum (Invitrogen) for 1 h at room temperature. The remaining immunostaining procedure was performed as previously described [65]. Mouse monoclonal anti-histone H4 (1:200; Abcam ab31830), rabbit polyclonal anti-H3K27Ac (1:100; Abcam ab4729), mouse monoclonal anti-Lam [97] (ADL84, 1:150), rabbit polyclonal anti-Elys [65] (1:3000) and guinea pig polyclonal anti-LBR [99] (1:1000) antibodies were applied. As the secondary, Alexa Fluor 546-conjugated goat anti-rabbit IgG (Invitrogen) or Alexa Fluor 488-conjugated goat anti-mouse IgG (Invitrogen), or Alexa Fluor 633-conjugated goat anti-guinea pig IgG (Invitrogen) antibodies were applied.

### *ImageJ* quantitation of chromatin distribution in the nucleus

Using *ImageJ*, fluorescence intensities of histone H4, H3K27Ac, Lam and LBR staining across the nucleus diameter of the equatorial focal plane of nuclei from Elys-KD or control S2 cells were extracted. Individual profiles were first normalized on the average intensity, then on the diameter of nucleus (delimited by peaks of LBR, or Lam fluorescence) and further aligned to determine the averaged profile. Data were obtained for 50 or 170 nuclei from Elys-KD or control S2 cells (25-100 nuclei per replicate) (S5 Table).

### Co-immunoprecipitation

Approximately 10^7^ S2 cells per sample were washed in PBS and homogenized in lysis buffer (50 mM Tris-HCl pH 7.5, 150 mM NaCl, 1 mM EDTA, 1% Triton X-100, 0.1% NP-40 and Complete Protease Inhibitor Cocktail (Roche)). The lysate was incubated for 20 min on ice prior to clearing by centrifugation at 16000 × g for 10 min at 4°C. Inputs were taken from the supernatant. 5 μl of rabbit polyclonal anti-Nup107 [98], 2 μl of rabbit polyclonal anti-Elys [65], or 4 μl of rabbit polyclonal anti-GAF [101] antibodies per sample were immobilized on Protein G Dynabeads (Thermo Fisher Scientific), incubated with lysates for 90 min at 25°C, after that beads were washed three times with washing buffer (PBS, 0.1% Tween-20 and 0.3% Triton X-100). The bound proteins were eluted by heating at 95°C for 10 min in the loading buffer containing SDS and 4 M urea. Control immunoprecipitation experiments using rabbit non-immune serum were performed in parallel.

### Hi-C in S2 cells upon Elys-KD

Hi-C experiments were performed in two biological replicates as described previously [102], with minor modifications to the fixation and lysis steps. ∼10^7^ control and Elys-KD S2 cells per replicate were fixed in PBS containing 2% formaldehyde for 10 min with occasional mixing. The reaction was quenched by the addition of 2 M glycine to a final concentration of 125 mM. Cells were lysed in 1.5 ml isotonic buffer [50 mM Tris−HCl pH 8.0, 150 mM NaCl, 0.5% (v/v) NP-40 substitute, 1% (v/v) Triton X-100, 1 × Halt™ Protease Inhibitor Cocktail (Thermo Scientific)] on ice for 15 min. Cells were pelleted by centrifugation at 2500 × g for 5 min, resuspended in 100 μl DpnII buffer (New England Biolabs), and pelleted again. All downstream steps were performed as described previously [102]. Hi-C libraries were sequenced on the Illumina NovaSeq 6000 at the Evrogen facility (www.evrogen.ru) resulting in 71–98 million 150-nt paired-end reads per sample (S10 Table).

### RNA-seq in S2 cells upon Elys-KD

∼10^7^ control and Elys-KD S2 cells were collected, and total RNAs were isolated using Trizol reagent (Invitrogen). Further steps of RNA-seq, such as isolation of poly(A^+^) RNAs on the oligo(dT) columns, synthesis of cDNAs using random primers, and sequencing of cDNA libraries resulting in 77–88 million 150-nt paired-end reads per sample (S10 Table), were performed at the GENEWIZ facility (www.genewiz.com).

### Analysis of DamID-seq data

Sequencing reads from two biological replicates of Dam, Dam-Lam, or Dam-Elys samples were adapter clipped and uniquely mapped to the dm3/R5 genomic assembly by *bowtie2* [103]. Reads were counted by *HTSeq-count* software [104] in the 0.3-kb genomic bins. Read counts were merged between replicates, as they were highly correlated (S12A Fig). The resulting read counts of Dam, Dam-Lam, or Dam-Elys samples were converted to reads per million mapped reads (RPM), normalized to RPM values of the Dam samples and log_2_ transformed. Next, Lam and Elys domain calling was performed with the two-state or three-state HMM algorithm, respectively (S2 and S3 Tables) (the scripts are deposited at GitHub: https://github.com/foriin/Elys_embryos_2022 and at Zenodo: https://zenodo.org/record/7808900#.ZDBCLvZBxPY). The log_2_ transformed profiles were visualized in UCSC genome browser using an auto-scale to data view and a smoothing window of 3 pixels.

### Analysis of external data

Various datasets were retrieved from GEO NCBI: Nup98_NPC and Nup98_nucl DamID profiles in Kc167 cells – from GSE19307 [28]; Lam DamID profile in Kc167 cells – from GSE20311 [5]; Lam DamID profiles in the central brain, Elav-positive neurons, Repo-positive glia and the fat body – from GSE109495 [6]; H3K27Ac ChIP-chip profile in S2 cells – from GSE20779 [74]; GAGA factor (GAF) ChIP-seq profile in S2 cells – from GSE40646 [105]; Nup98, Elys and Nup93 ChIP-seq profiles and peaks in S2 cells – from GSE94922 [35] and GSE135610 [37]; STARR-seq enhancer profiles in S2 cells – from GSE40739 [83] and GSE47691 [84]. The enrichment regions for Nup98_NPC, Nup98_nucl, Lam, H3K27Ac and STARR-seq enhancer profiles were identified with two-state (for Lam and STARR-seq enhancers) or three-state (for Nup98_NPC, Nup98_nucl and H3K27Ac) HMM. Less than 900-bp gaps between domains were filled in. For peak calling of ChIP-seq profiles, *MACS2* [106] was applied. Domain/domain or gene/domain intersections were computed in *R* as a ratio of genome coverage using the *GenomicRanges* package in *Bioconductor* [107]. To perform permutation analysis we invoked *BEDTools* [108] in *R* to reshuffle 10000 times various sites of Elys, Nup98, genes, or enhancers, counted the number of intersected sites and compared them with the original data.

### Analysis of RNA-seq data

Sequencing reads from two biological replicates of control and Elys-KD S2 cells were adapter clipped. Low-quality reads with the length of less than 20 nt were fltered out and the remaining reads were uniquely mapped to dm3/R5 genome assembly. To test whether RNA-seq replicates were similar, sequencing reads were counted per 0.3-kb genomic bins. Then, counts were normalized for sequencing depth and clustering analysis was performed (S12B Fig). Since replicates were highly correlated, they were merged. Reads were counted for genes of FlyBase r5.57 annotation and converted to transcripts per million (TPM) values using the *Salmon* tool [109] (S8 Table). An analysis of resulting data was performed in *RStudio IDE* using the *R* packages *tximport*, *GenomicRanges*, *dplyr* and *ggplot2*. Differentially expressed genes upon Elys-KD were identifed by the *DESeq2* package [110] for *R* with the cutoff parameters *P* < 0.05 and TPM FC > 1.5 (S8 Table).

### Analysis of Hi-C data

Sequencing reads from two biological replicates of Hi-C control and Elys-KD experiments were trimmed with *Trim Galore* and processed with the *distiller-nf* pipeline (version 0.3.3) (https://doi.org/10.5281/zenodo.3350926). Parameter “*stringency 3*” was applied during the trimming procedure. Reads were mapped to chromosomes 2L, 2R, 3L, 3R, 4, X, and M of the dm3/R5 *Drosophila* genome assembly with the default settings and with an option *MAPQ_30* (mapq1>=30 and mapq2>=30). For the downstream analysis of the interaction matrix, contacts between genomic bins separated by < 1 kb were removed. After that, only chromosomes 2L, 2R, 3L, 3R, and X were considered. For the replicate comparison, Hi-C maps were downsampled with the *cooltools* package (https://cooltools.readthedocs.io/en/latest/) to nearly the same total number of contacts across all replicates. Then, the iterative correction (IC) [111] was applied, and samples were clustered with the *R* package *HiCRep* [112] based on the stratum-adjusted correlation coefficient (SCC). Clustering was performed on 4-kb resolution Hi-C maps for bins separated by ≤ 1 Mb. SCCs obtained for each chromosome were averaged and used to build a hierarchical dendrogram (S12C Fig). Next, replicates were merged with the *cooler* package (https://cooler.readthedocs.io/en/latest/), downsampled to nearly the same total counts, and corrected by IC, resulting in the final Hi-C maps.

TAD boundaries were called with the *Armatus* (version 2.2) software [78] (https://github.com/kingsfordgroup/armatus) in 4-kb resolution Hi-C maps, which were additionally preprocessed. Each interchromosomal matrix was linearly interpolated, the contacts exceeding the range between the 1^st^ and the 99^th^ percentiles were replaced with the corresponding values of these percentiles and, finally, matrices were ln-transformed. Then *Armatus* was run for autosomes with the parameter *γ* = 1. For the X chromosome, *γ* = 0.6 and *γ* = 0.4 were chosen for control and Elys-KD maps respectively. For the downstream analysis, TADs were divided into 3 groups by their activity. To simultaneously account for the LAD coverage and the proportion of active chromatin (states 1 and 2 [74]), the Jaccard coefficient between these two metrics was calculated by the formula: Jacc = (X-Y)/(X+Y), where X is the proportion of LADs and Y is the proportion of chromatin states 1 and 2 in the TAD. The most active TADs with Jaccard coefficient < -0.8 were attributed to group A. TADs of intermediate activity with Jaccard coefficient ≥ -0.8 and ≤ 0.8 were attributed to group B. Finally, inactive TADs with Jaccard coefficient > 0.8 were attributed to group C. ACF was calculated based on the distance-normalized (observed/expected) IC Hi-C maps as the mean value of contacts within a TAD.

Active (A) and inactive (B) compartments [77] were identified for 10-kb resolution IC Hi-C maps with the *cooltools* package. Positive values of the first principal component (PC1) corresponding to the A compartment were selected by correlation with gene expression. *Cooltools* package was used for the saddle plot generation, which included the following steps. For each chromosome, bins were ranked according to their PC1 values and 1% of bins with the highest and lowest PC1 values were filtered out. Then, distance-normalized contact frequencies were averaged within quantiles of the PC1, and the resulting values were used for generation of saddle plots. We applied the same number of PC1 quantiles for both A and B compartments across all chromosomes and experimental conditions to analyze changes of compartmentalization, averaged by chromosomes. Box-plots showing the saddle-plot difference between Elys-KD and control include interactions between the bins falling within the 10 highest and lowest quantiles of PC1.

The insulation score (IS) [113] was calculated for 2-kb resolution Hi-C maps. The modified IS (IS^m^) was calculated as the average distance-normalized contact frequency within the 8-kb square, starting at the second diagonal. Thus, contact frequency within a bin containing Elys_NPC site and its interactions with the adjacent bins were not taken into account. Averaged IS^m^-curves were smoothed using linear interpolation.

### Generation of averaged profiles

Averaged profiles and A/T profiles were generated in *RStudio* using the *R* packages *GenomicRanges*, *dplyr ggplot2, Biostrings, seqinr* and *rtracklayer*. Metagene profiles were generated using *deepTools2* software [114] and *RStudio*. Heatmaps of Lam profiles were generated using *R* with *genomation* package [115]. Custom *R* package for working with *D. melanogaster* dm3 genome release can be found at https://github.com/foriin/dm3.

### DNA sequence motif analysis

For sequence motif identification, *MEME 5.3.3* with the default settings was applied [75].

### Statistical analysis

The Mann-Whitney (M-W) U-test was applied for *P*-values estimation upon comparison of two distributions. The Wilcoxon signed-rank test was applied for estimation of whether log_2_ FC values was symmetric around zero. *P*-values for occasional gene/site or site/site overlapping were estimated by permutation test with 10000 permutations.

## Data availability

Raw and processed DamID-seq, RNA-seq, and Hi-C data were deposited in the NCBI Gene Expression Omnibus (GEO) under the accession numbers GSE219152 (for DamID-seq, RNA-seq) and GSE218886 (for Hi-C). In-house scripts are available at GitHub (https://github.com/foriin/Elys_embryos_2022) and at Zenodo: (https://zenodo.org/record/7808900#.ZDBCLvZBxPY). The link for UCSC Browser (http://hgw1.soe.ucsc.edu/cgi-bin/hgTracks?db=dm3&lastVirtModeType=default&lastVirtModeExtraState=&virtModeType=default&virtMode=0&nonVirtPosition=&position=chr2R%3A16166667%2D16833333&hgsid=1515860147_HDWc9AODZV9tmQHWMmf4XWu4Wfx5).

## Funding

This work was supported by the Russian Ministry of Science and Higher Education (075-15-2021-1062). Hi-C experiments were supported by the Russian Science Foundation (21-64-00001 to SVR). Hi-C data analysis was supported by the Russian Foundation for Basic Research (21-34-70051 to EEK). The funders had no role in study design, data collection and analysis, decision to publish, or preparation of the manuscript.

## Competing interests

The authors have no competing interests.

## Acknowledgements

We thank Paul Fisher for anti-Lam antibodies, Georg Krohne for anti-LBR antibodies, Gary Karpen for anti-CenpA antibodies, Valerie Doye for anti-Nup107 antibodies, Maksim Erokhin for anti-GAF antibodies, Centre of Common Scientific Equipment of the National Research Centre “Kurchatov Institute” for providing access to confocal microscope.

## Author contributions

YYS conceived the project. SAD generated the construct for DamID and carried out DamID in embryos, performed RNAi experiments, Western-blot analysis, co-immunoprecipitation, immunostaining, TUNEL and FISH experiments. MAS (supervised by SVU) performed Hi-C experiments. EAM maintained cell cultures and performed some immunostaining. OMO performed fly crossing. SAL performed immunostaining of polytene chromosomes. AAF carried out early FISH experiments. AYI carried out FISH experiments with *36C* probe. AAI (supervised by YYS) analyzed various data sets (DamID, RNA-seq and external data). ADK (supervised by EEK) analyzed Hi-C data. VVN analyzed FISH results. YYS wrote the draft of the manuscript and prepared figures. SVU, EEK, and SVR made significant contribution to the final version with the input of all other authors.

## Supporting information

**S1 Fig. An efficiency of Elys-KD.** (**A**–**D**) Western-blot analysis of protein extracts from control, Elys-KD, LamKD or Elys-KD + Lam-KD S2 cells probed by anti-Elys, anti-Lam or anti-Actin (loading control) antibodies. Quantification of depletion efficiency was performed in *ImageJ*. In (**A**–**C**) protein extracts from two replicates were combined before loading.

**S2 Fig. Upon Elys-KD, mRNA export is not impaired.** (**A**) Confocal images of control or Elys-KD S2 cells after oligo(dT) FISH (violet) and immunostaining with anti-Lam (green) and anti-Elys (red) antibodies. Scale bar 1 µm. (**B**) Box-plots showing ratios of fluorescence intensities (in cytoplasm to nucleus) for oligo(dT) FISH. Fluorescence in the 40 control and Elys-KD cells (from two replicates) was quantified using *ImageJ*. *P* value was estimated in a M-W U-test. n.s. - non-significant (*P* > 0.05).

**S3 Fig. TUNEL assay did not detect an increase in the proportion of apoptotic cells upon Elys-KD.** (**A**) Confocal image of control or Elys-KD S2 cells stained with TUNEL assay (violet), as well as with anti-Elys (green) and anti-Lam (blue) antibodies. Scale bar 30 µm. (**B**) Box-plots showing percentage of apoptotic cells in control or Elys-KD S2 cells. *P* value was calculated in a M-W U-test. n.s. -non-significant (*P* > 0.05).

**S4 Fig. Elys_NPC and Elys_nucl sites mostly overlap with inactive and active chromatin types, respectively.** (**A**,**B**) Pie charts showing percentage of overlap between Elys_NPC, Elys_nucl or Elys_NPC/nucl sites with chromatin domains identified according to 5-state chromatin model in Kc167 cells (**A**), or according to 9-state chromatin model in S2 cells (**B**).

**S5 Fig. Weak dips in Lam profile at positions of Elys ChIP-seq peaks from S2 cells.** (**A**–**C**) Averaged Lam_Kc167, Lam_embryo, Elys_embryo, Elys_S2 and H3K27Ac profiles centered at randomly chosen positioned within Kc167 LADs, or centered at Elys_NPC sites located within Kc167 LADs overlapped with inactive chromatin (states 6-9), or centered at Elys_S2 ChIP-seq sites located within Kc167 LADs overlapped with inactive chromatin (states 6-9) (**A**), or centered at Elys_S2 ChIP-seq sites located within Kc167 or embryo LADs (±2 kb from LAD boundaries) (**B**), or centered at Nup93_S2 ChIP-seq sites (**C**).

**S6 Fig. The dips in Lam profile are revealed at positions of Elys_NPC sites during cell differentiation.** (**A**) Screenshot from UCSC genome browser showing H3K27Ac (orange), Nup98_nucl (pink), Nup98_NPC (violet), Elys_embryo (red), Elys_S2 (yellow), Lam_embryo and Lam_Kc167 (brown) profiles, as well as Lam_brain, Lam_neurons, Lam_glia and Lam_fat body (black) profiles for the representative region of chromosome 3R. The corresponding domains are provided as the rectangles over profiles. 9-state chromatin model, RNA-seq in control S2 cells and RefSeq genes are indicated below. The overlapped regions between Elys_embryo and Nup98_NPC domains containing Elys_NPC peaks are outlined by translucent rectangles (they mostly coincide with the dips in Lam profiles during cell differentiation). (**B**) Averaged Lam_brain, Lam_neurons, Lam_glia and Lam_fat body profiles centered at Elys_NPC sites, located within LADs, from the corresponding cell type. (**C**) Heatmaps of Lam profiles centered at Elys_NPC sites, located within LADs, from the corresponding cell type sorted from minimal (top) to maximal (bottom) values at the central bins. Rows where central bins had zero value either in Dam-Lam or in Dam profiles were removed. Bins with zero value of either Dam-Lam or Dam in other locations are marked by grey color.

**S7 Fig. *60D*, *22A* and *36C* hybridization probes and/or their nearby regions contain Elys_NPC sites.** (**A**–**C**) Screenshots from UCSC genome browser showing H3K27Ac (orange), Nup98_nucl (pink), Nup98_NPC (violet), Elys_embryo (red), Lam_Kc167 and Lam_embryo (brown) profiles, as well as the corresponding domains (rectangles over profiles) for the *60D* (**A**), *22A* (**B**), and *36C* (**C**) regions. RefSeq genes are indicated below. Hybridization probes are indicated by black rectangles on the top of the panels. Exact boundaries of the **60D** probe are unknown.

**S8 Fig. Analysis of chromatin distribution in nuclei upon Elys-KD.** (**A**) Staining of control and Elys-KD cells with anti-LBR (violet), anti-histone H4 (green) and anti-Elys (red) antibodies. (**B**) Staining of control and Elys-KD cells with anti-Lam (pink), anti-H3K27Ac (green) and anti-Elys (red) antibodies. Scale bars 10 µm.

**S9 Fig. Active chromatin became more compact, whereas inactive - less compact upon loss of Elys in S2 cells.** (**A**,**B**) Log_2_ fold-change (FC) of ACF (Elys-KD/control) in all TADs (**A**) or in the X chromosomal TADs (**B**) ranked by quartiles of total gene expression within them (according to RNA-seq data (in RPM) in control S2 cells, where 1^st^ quartile corresponds to the lowest, and 4^th^ quartile - to the highest gene expression). *P*-values were estimated in a Wilcoxon signed-rank test. (**C**) Saddle plots showing values of aggregated contact frequency for the intra- chromosomal contacts in autosomes in control (left panel) and Elys-KD (right panel) cells ranked by PC1 values. (**D**) Box-plots showing subtraction of aggregated contact frequency (Elys-KD minus control) log_2_FC(observed/expected) for only the X chromosome within active (AA), inactive (BB), and between active and inactive (AB) chromatin compartments. *P*-values were estimated in a Wilcoxon signed-rank test.

**S10 Fig. Pie chart showing percentage of Elys binding sites overlapped with genes or intergenic regions.**

**S11 Fig. Elys is enriched at the 5’- and 3’-ends of genes.** (**A**–**C**) Metagene profiles for Elys (red) and A/T content (blue) (**A**), Elys (red) and H3K27Ac (blue) (**B**), Elys (red) and GAF (blue) (**C**) over genes containing Elys_nucl, Elys_NPC sites or combinations of these sites. (**D**) Western-blot analysis does not reveal co-immunoprecipitation of Elys and GAF. Westerns are stained with anti-Elys (left panel) or anti-GAF (right panel) antibodies. Anti-GAF antibodies detect two GAF isoforms. IP/input ratio 1:4.5.

**S12 Fig. Replicates clustering.** Replicates for DamID-seq (**A**), RNA-seq (**B**) and Hi-C (**C**) are highly correlated according to the Spearman correlation coefficient (**A**,**B**), or to the stratum-adjusted correlation coefficient (SCC) (**C**).

**S1 Table. TUNEL assay analysis.**

**S2 Table. Elys_embryo, Elys_NPC, Elys_nucl and Elys_NPC/nucl sites.**

**S3 Table. LADs_embryo.**

**S4 Table. FISH analysis.**

**S5 Table. Distribution of histone H4 or H3K27Ac fluorescence intensity across a nucleus.**

**S6 Table. Volume of nuclei.**

**S7 Table. TADs.**

**S8 Table. RNA-seq analysis.**

**S9 Table. Primers for PCR amplification. S10 Table. NGS statistics.**

## Notes

### Competing Interest Statement

The authors have declared no competing interest.

