## Supplementary Figures for "Nucleoporin Elys attaches peripheral chromatin to the nuclear pores in interphase nuclei"

S1 Fig

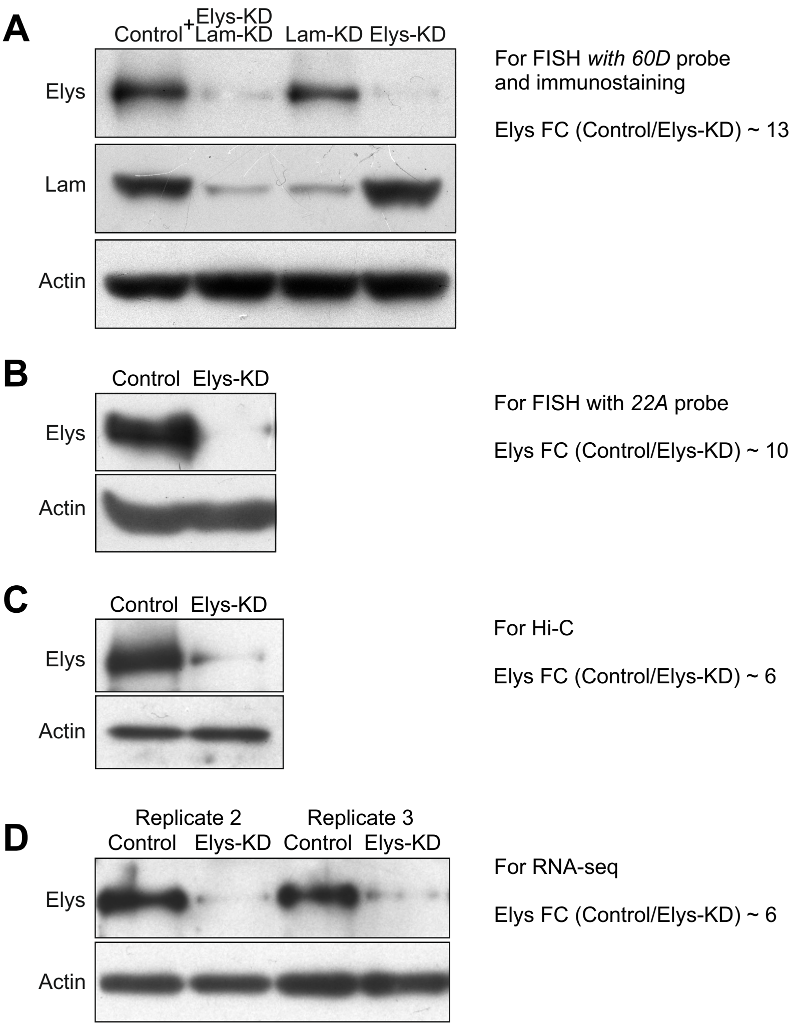
A
Control
Elys-KD
+
Lam-KD
Lam-KD
Elys-KD
Elys
Lam
Actin
For FISH with 60D probe
and immunostaining
Elys FC (Control/Elys-KD) ~ 13
B
Control
Elys-KD
Elys
Actin
For FISH with 22A probe
Elys FC (Control/Elys-KD) ~ 10
C
Control
Elys-KD
Elys
Actin
For Hi-C
Elys FC (Control/Elys-KD) ~ 6
D
Replicate 2
Replicate 3
Control
Elys-KD
Control
Elys-KD
Elys
Actin
For RNA-seq
Elys FC (Control/Elys-KD) ~ 6

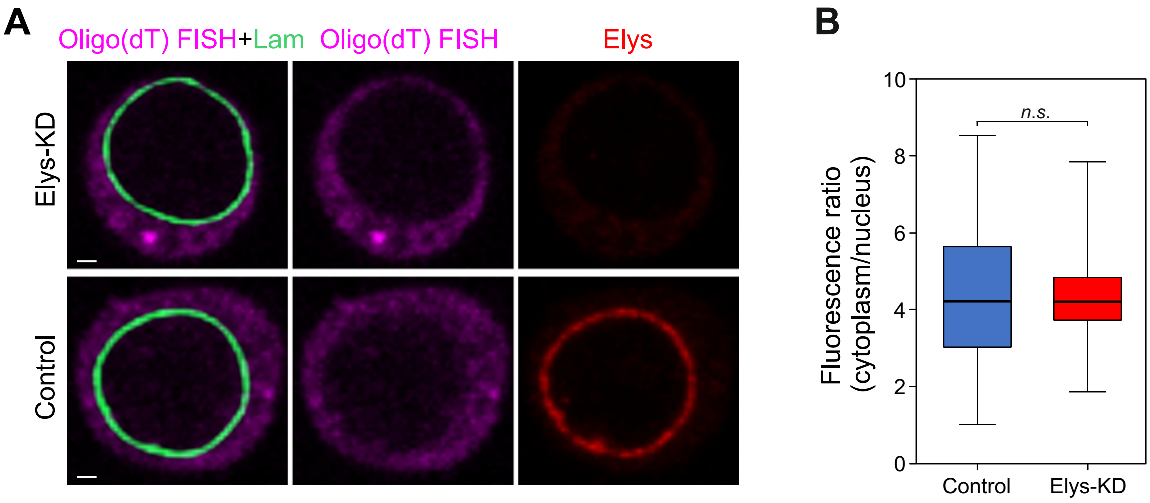
A
Oligo(dT) FISH+Lam
Oligo(dT) FISH
Elys
Elys-KD
Control
B
Fluorescence ratio (cytoplasm/nucleus)
n.s.
0
2
4
6
8
10
Control
Elys-KD

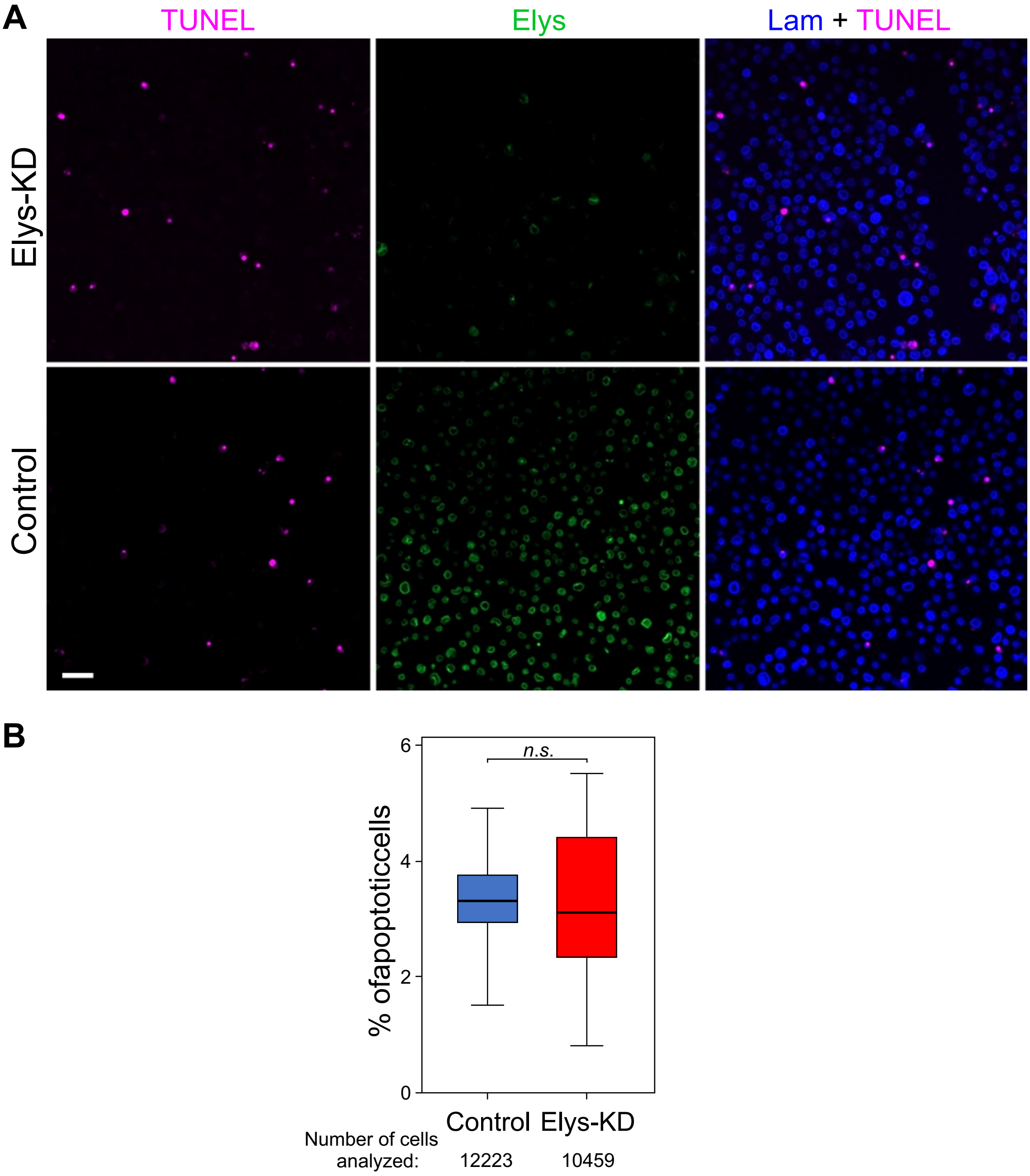

A
TUNEL
Elys
Lam + TUNEL
Elys-KD
Control
B
n.s.
% ofapoptoticcells
0
2
4
6
Control
Elys-KD
Number of cells
analyzed:
12223
10459

S4 Fig

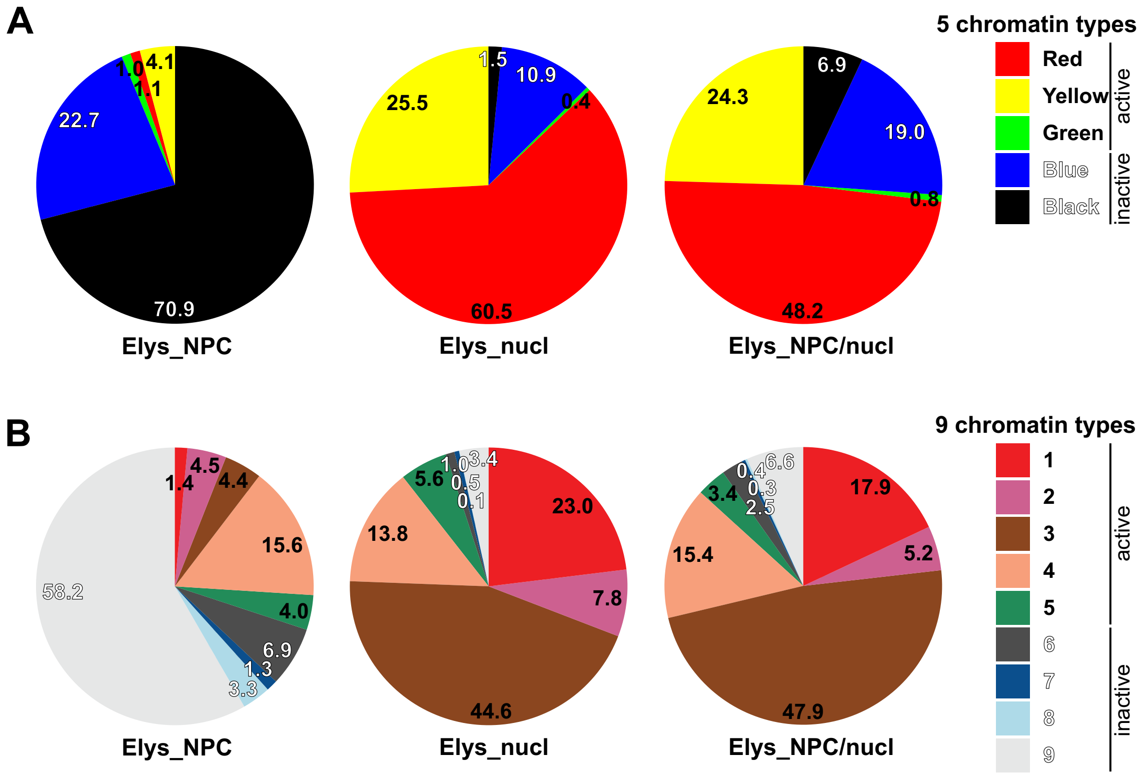
A
5 chromatin types
Red
Yellow
Green
Blue
Black
active
inactive
4.1
1.0
1.1
22.7
70.9
Elys_NPC
1.5
10.9
0.4
25.5
60.5
Elys_nucl
6.9
24.3
19.0
0.8
48.2
Elys_NPC/nucl
B
9 chromatin types
1
2
3
4
5
6
7
8
9
active
inactive
1.4
4.5
4.4
15.6
4.0
6.9
1.3
3.3
58.2
Elys_NPC
1.0
3.4
0.5
0.1
5.6
23.0
13.8
7.8
44.6
Elys_nucl
0.4
6.6
0.3
2.5
3.4
17.9
15.4
5.2
47.9
Elys_NPC/nucl

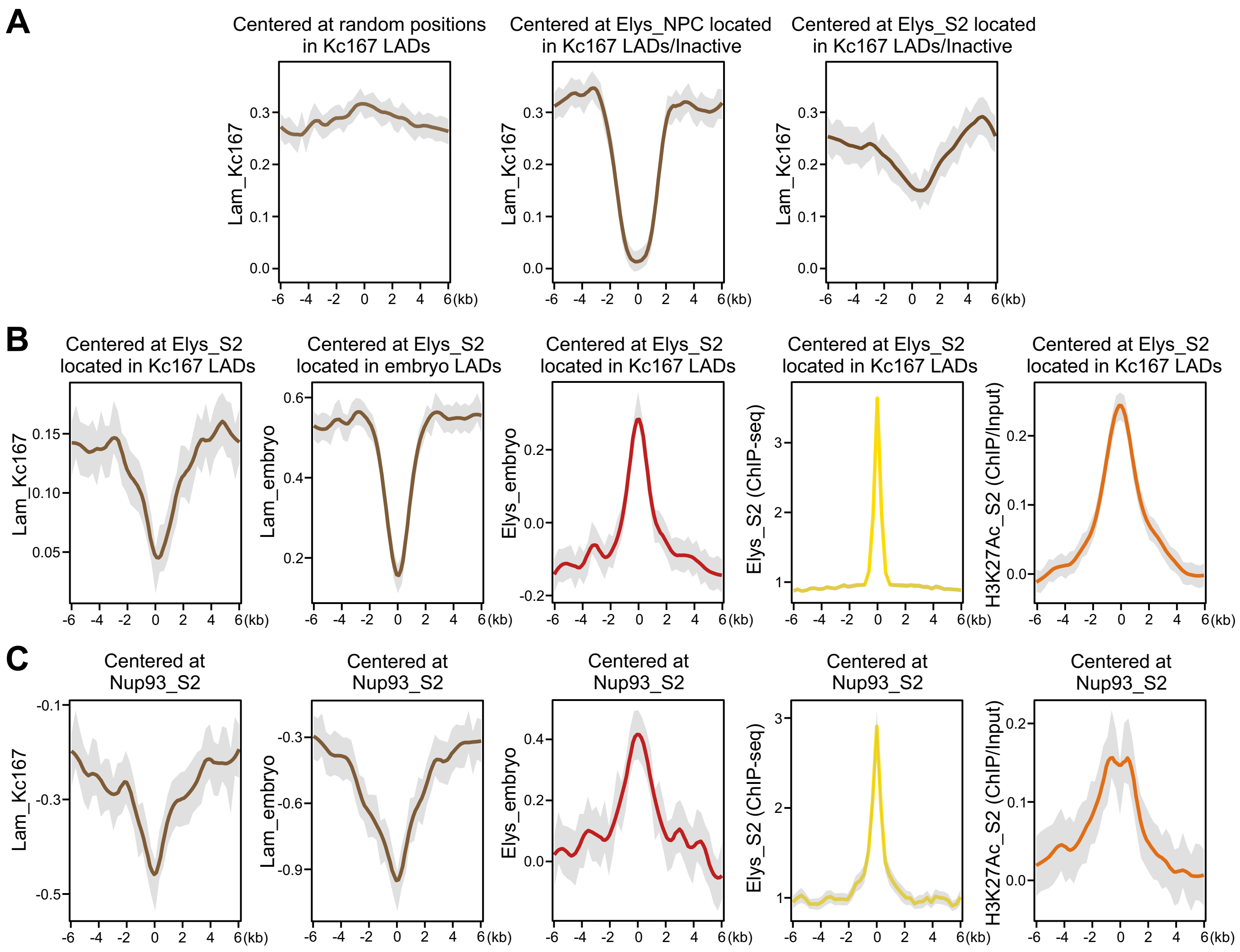
A
Centered at random positions in Kc167 LADs
Centered at Elys_NPC located in Kc167 LADs/Inactive
Centered at Elys_S2 located in Kc167 LADs/Inactive
Lam_Kc167
(kb)
B
Centered at Elys_S2 located in Kc167 LADs
Centered at Elys_S2 located in embryo LADs
Lam_Kc167
Lam_embryo
Elys_embryo
Elys_S2 (ChIP-seq)
H3K27Ac_S2 (ChIP/Input)
C
Centered at Nup93_S2

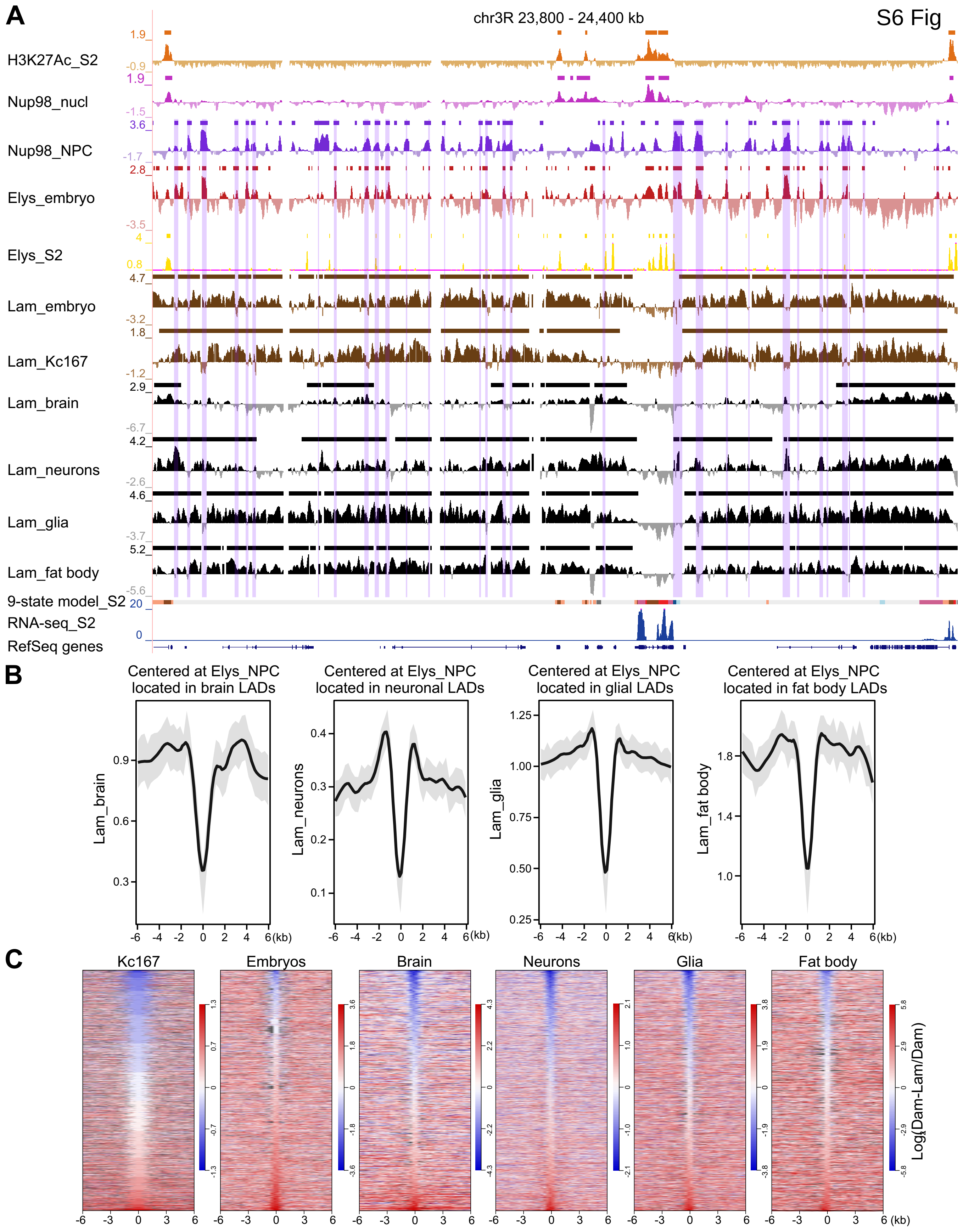


locations are marked by grey color.

### S7 Fig

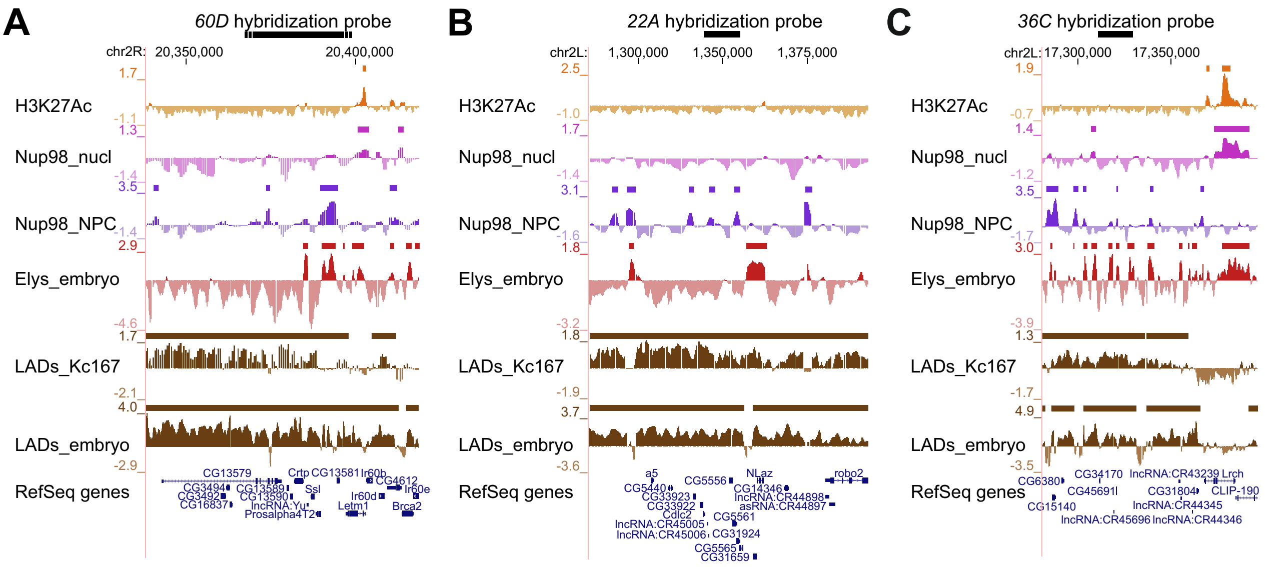
A
60D hybridization probe
chr2R: 20,350,000
20,400,000
H3K27Ac
1.7
-1.1
Nup98_nucl
1.3
-1.4
Nup98_NPC
3.5
-1.4
Elys_embryo
2.9
-4.6
LADs_Kc167
1.7
-2.1
LADs_embryo
4.0
-2.9
RefSeq genes
B
22A hybridization probe
chr2L: 1,300,000
1,350,000
1,375,000
H3K27Ac
2.5
-1.0
Nup98_nucl
1.7
-1.4
Nup98_NPC
3.1
-1.6
Elys_embryo
1.8
-3.2
LADs_Kc167
1.8
-1.9
LADs_embryo
3.7
-3.6
RefSeq genes
C
36C hybridization probe
chr2L: 17,300,000
17,350,000
H3K27Ac
1.9
-0.7
Nup98_nucl
1.4
-1.2
Nup98_NPC
3.5
-1.7
Elys_embryo
3.0
-3.9
LADs_Kc167
1.3
-1.7
LADs_embryo
4.9
-3.5
RefSeq genes

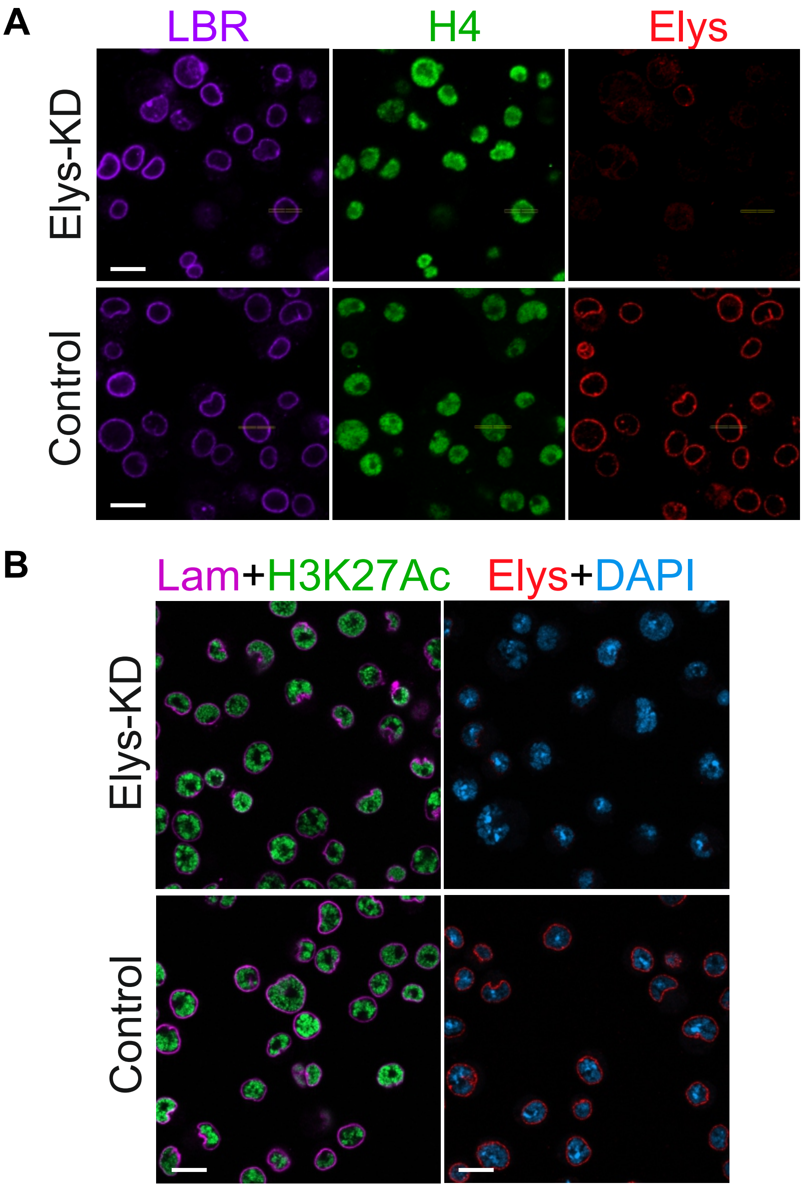
A
LBR
H4
Elys
Elys-KD
Control
B
Lam+H3K27Ac
Elys+DAPI
Elys-KD
Control

S9 Fig

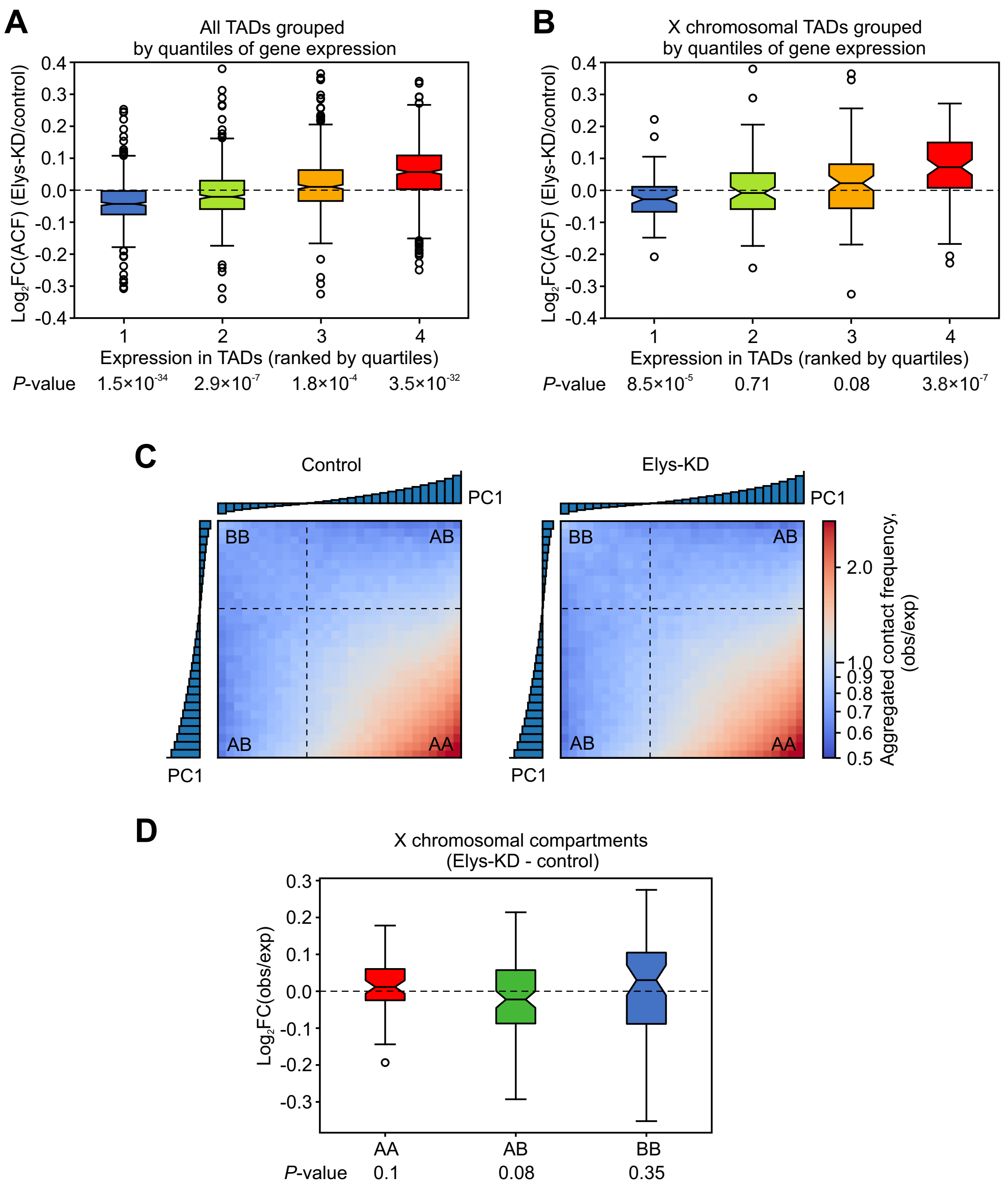
A
All TADs grouped
by quantiles of gene expression
Log2FC(ACF) (Elys-KD/control)
0.4
0.3
0.2
0.1
0.0
-0.1
-0.2
-0.3
-0.4
1
2
3
4
Expression in TADs (ranked by quartiles)
P-value
1.5×10-34
2.9×10-7
1.8×10-4
3.5×10-32
B
X chromosomal TADs grouped
by quantiles of gene expression
Log2FC(ACF) (Elys-KD/control)
Expression in TADs (ranked by quartiles)
P-value
8.5×10-5
0.71
0.08
3.8×10-7
C
Control
Elys-KD
PC1
BB
AB
AA
Aggregated contact frequency,
(obs/exp)
2.0
1.0
0.9
0.8
0.7
0.6
0.5
D
X chromosomal compartments
(Elys-KD - control)
Log2FC(obs/exp)
AA
AB
BB
P-value
0.1
0.08
0.35

S10 Fig

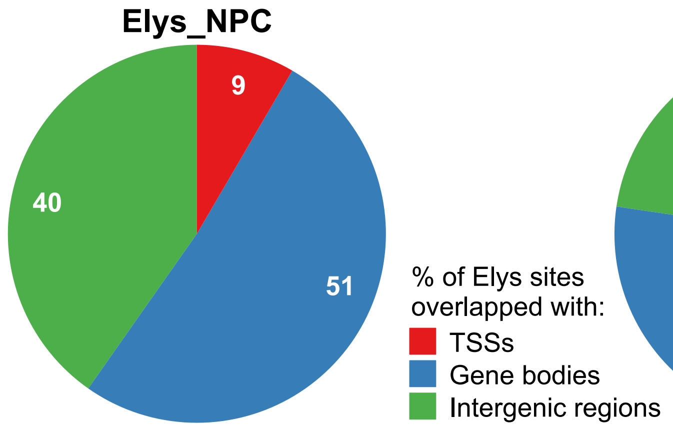

Elys_NPC
9
51
40
% of Elys sites
overlapped with:
TSSs
Gene bodies
Intergenic regions
Elys_nucl
42
35
23

S11 Fig

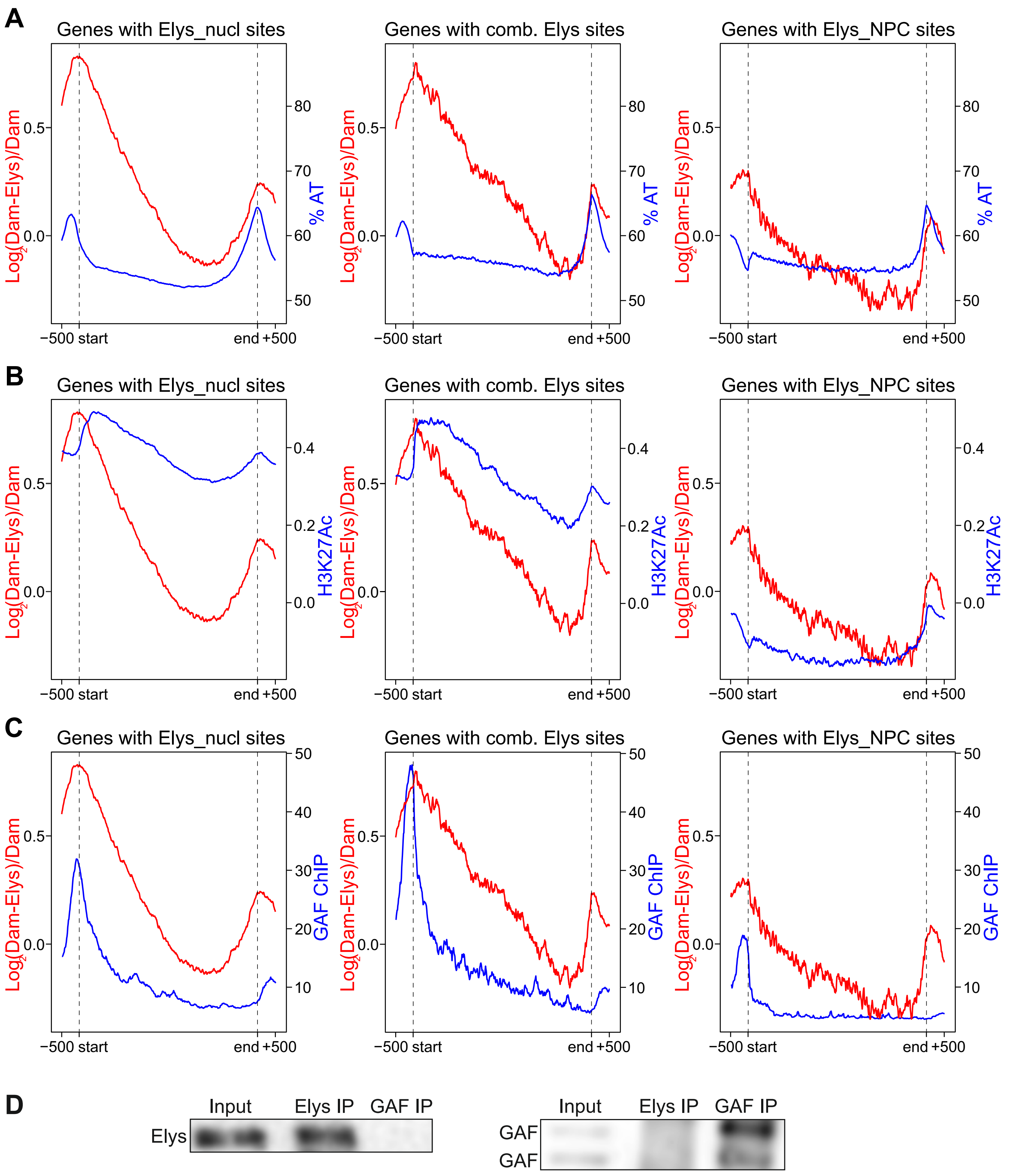
A
Genes with Elys_nucl sites
Genes with comb. Elys sites
Genes with Elys_NPC sites
Log2(Dam-Elys)/Dam
% AT
0.0
0.5
50
60
70
80
−500 start
end +500
B
Genes with Elys_nucl sites
Genes with comb. Elys sites
Genes with Elys_NPC sites
Log2(Dam-Elys)/Dam
H3K27Ac
0.0
0.2
0.4
0.5
−500 start
end +500
C
Genes with Elys_nucl sites
Genes with comb. Elys sites
Genes with Elys_NPC sites
Log2(Dam-Elys)/Dam
GAF ChIP
0.0
0.5
10
20
30
40
50
−500 start
end +500
D
Input
Elys IP
GAF IP
Elys
GAF
GAF

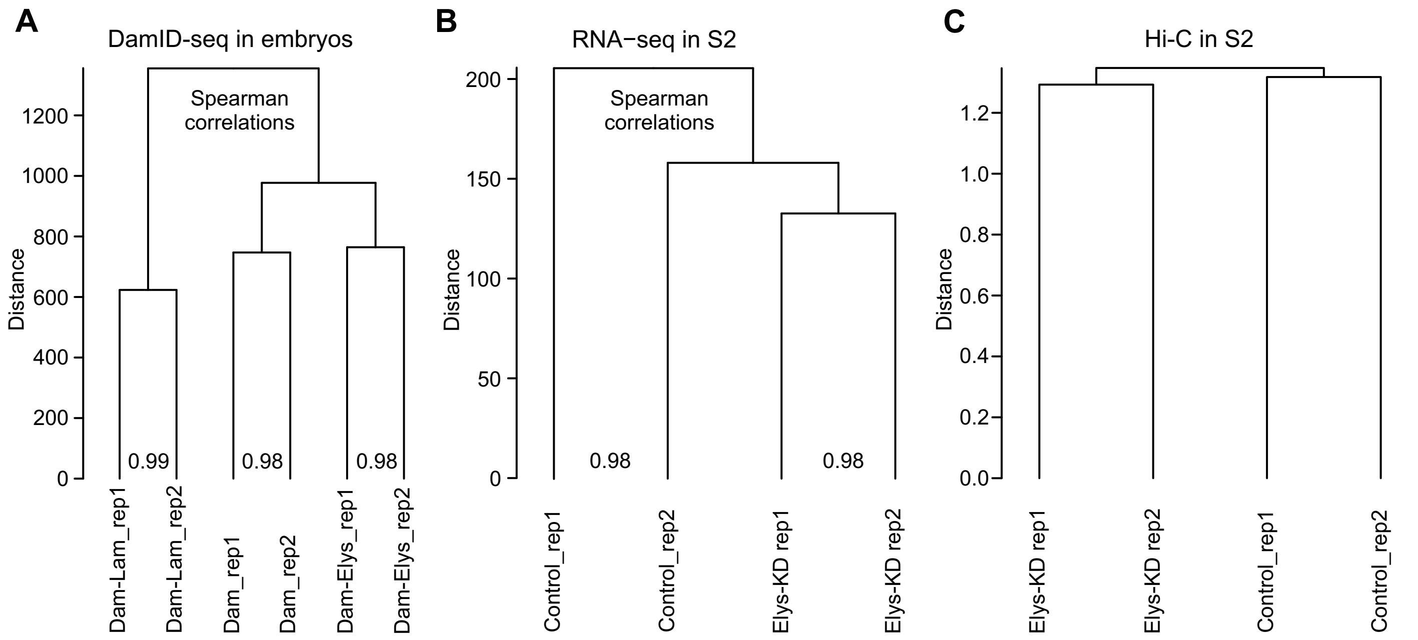
A
DamID-seq in embryos
Spearman
correlations
Distance
0
200
400
600
800
1000
1200
0.99
0.98
0.98
Dam-Lam_rep1
Dam-Lam_rep2
Dam_rep1
Dam_rep2
Dam-Elys_rep1
Dam-Elys_rep2
B
RNA−seq in S2
Spearman
correlations
Distance
0
50
100
150
200
0.98
0.98
Control_rep1
Control_rep2
Elys-KD rep1
Elys-KD rep2
C
Hi-C in S2
Distance
0.0
0.2
0.4
0.6
0.8
1.0
1.2
Elys-KD rep1
Elys-KD rep2
Control_rep1
Control_rep2
